# Treated like dirt: Robust forensic and ecological inferences from soil eDNA after challenging sample storage

**DOI:** 10.1101/2021.12.22.473824

**Authors:** Tobias Guldberg Frøslev, Rasmus Ejrnæs, Anders J. Hansen, Hans Henrik Bruun, Ida Broman Nielsen, Flemming Ekelund, Mette Vestergård, Rasmus Kjøller

**Affiliations:** Section for Geogenetics, GLOBE Institute, University of Copenhagen, Øster Voldgade 5-7, 1350 Copenhagen, Denmark; Section for Biodiversity & Conservation, Department of Ecoscience, Aarhus University, Grenåvej 14, 8410 Rønde, Denmark; Department of Biology, University of Copenhagen, Universitetsparken 15, 2100 Copenhagen, Denmark; Section for Evolutionary Genomics, GLOBE Institute, Øster Farimagsgade 5, 1353 Copenhagen, Denmark; Section for Entomology and Plant Pathology, Department of Agroecology, Aarhus University, 4200 Slagelse, Denmark

## Abstract

Biodiversity of soil microbiota is routinely assessed with environmental DNA-based methods, among which amplification and massive parallel sequencing of marker genes (eDNA metabarcoding) is the most common. Soil microbiota may for example be investigated in relation to biodiversity research or as a tool in forensic investigations.

After sampling, the taxonomic composition of soil biotic communities may change. In order to minimize community changes after sampling, it is desirable to reduce biological activity, e.g. by freezing immediately after sampling. However, this may be impossible due to remoteness of study sites or, in forensic cases, where soil has been attached to a questioned item for protracted periods of time.

Here we investigated the effect of storage duration and conditions on the assessment of the soil biota with eDNA metabarcoding. We extracted eDNA from freshly collected soil samples and again from the same samples after storage under contrasting temperature conditions.

We used five different primer sets targeting bacteria, fungi, protists (cercozoans), general eukaryotes, and plants. For these groups, we quantified differences in richness, evenness and community composition. Subsequently, we tested whether we could correctly infer habitat type and original sample identity after storage using a large reference dataset.

We found increased community composition differences with extended storage time and with higher storage temperature. However, for samples stored less than 28 days at a maximum of 20°C, changes were generally insignificant. Classification models could successfully assign most stored samples to their exact location of origin and correct habitat type even after weeks of storage. Even samples showing larger compositional changes generally retained the original sample as the best match (relative similarity).

Our results show that for most biodiversity and forensic applications, storage of samples for days and even several weeks may not be a problem, if storage temperature does not exceed 20°C. Even after suboptimal storage conditions, significant patterns can be reproduced.

## 1 Introduction

A teaspoon of soil may contain more than a billion bacterial cells, meters of fungal hyphae and profuse numbers of protists, nematodes and small arthropods^1^. Moreover, the phylogenetic diversity of soil is stunning not only at global scale, but also at local scales^2–5^. Today, high-throughput sequencing – often with the approach called eDNA metabarcoding – is the standard tool for mapping this enormous microbial biodiversity. DNA metabarcoding has shown that soil microbial biodiversity varies at scales from global to local, with a strong impact of habitat^6,7^. The high microbial diversity in combination with different habitat requirements for most microorganisms, make the microbial composition of any soil sample unique, and with a compositional signature that reflects the habitat and sampling location. The continuous introduction and extinction of microbial species to any specific site further contributes to the uniqueness of any point or snapshot sample of the soil community. This ecological fingerprint may be used for making inferences about the wider community surrounding the sampling location, of potential use in ecological studies ^8,9^, as well as in forensics^10^.

Almost a century ago, Edmond Locard stated that a perpetrator of a crime will bring something to the crime scene, and leave with something from it^11^. Soil is ubiquitous, and has thus been of forensic interest for a long time, as it has the potential of linking persons or objects to a crime scene^12^. Until recently biotic forensic soil analyses have been based solely on microscopic analyses. Hence, they have been restricted to a relatively small proportion of the actual biotic component, and dependent on the skills of a few highly trained experts^13^. High throughput sequencing extends the scope to all biotic components, introduces methods that can be standardized, and produces relatively objective data, which may easily be analyzed with common statistical approaches^14–16^.

Two basic types of forensic cases can be identified – matching and provenance prediction. In cases of matching (or discrimination), the likelihood that two soil samples share the same origin in space is assessed – e.g. soil from a suspect’s shoe sole and soil from a crime scene. Here, DNA metabarcoding has a huge potential^17–23^. Provenance prediction can be used, when no potential crime scenes have been identified. Here, the likely origin(s) of the questioned sample is narrowed down in terms of a potential geographical area or habitat/location type. Provenance prediction using soil DNA metabarcoding has so far only been explored in a single study^10^, but the same overall approach also has proven useful for dust samples^24^.

For biodiversity studies and forensic applications alike, it is important that the detected community of the sample reflects the biotic composition at the sampling site with a level of representativity adequate for the research question. For any soil sample, the final detected community will depend on its actual taxonomic composition, analytical bias and variance from the laboratory procedures, and finally the selected bioinformatic and statistical approaches. For eDNA metabarcoding, a number of sources of variance and errors relate to the last-mentioned points: e.g. DNA extraction method, PCR setup, sample tagging, library building approach, contamination, sequencing platform, and sequence processing/filtering, OTU definition, and statistical approaches^25–27^. These sources of variance/error mainly influence comparability between data from different studies, and to a great extent they can be controlled and standardized by the researcher. In contrast, what happens to a sample before it arrives in the lab may be less easy to control and standardize. Pre-analytical handling and storage are known to result in changes in the taxonomic composition, especially for heterotrophic microorganisms sensitive to the altered conditions.

To minimize biotic activity immediately after sampling, most sampling protocols prescribe to cool/freeze samples or add a buffer that inactivates biotic activity^28,29^. In forensic applications, a soil sample recovered from an object or from a suspect has usually been removed from the crime scene for days, weeks or months and therefore has been subjected to desiccation or temperatures different from its original conditions. These “sample storage” conditions potentially change the biotic composition of the sample, which will ultimately affect the interpretation of laboratory results. Thus, it is important to establish a range of storage times and conditions that allow a valid interpretation of the different biotic components of soil samples. A study investigating the storage effect on three different soils using a small set of realistic storage scenarios for biodiversity studies concluded that the different approaches only marginally impaired the inferred richness measures and community patterns^30^. They did find changes in richness, but the effect was insignificant, if rare taxa were not considered. In their proposed guidelines, they advocated storage at 4°C for shorter periods (if possible), and otherwise desiccation of the sample with silica gel. Another study examined the forensic application of bacterial soil communities with a set of samples and locations, mimicking realistic evidence samples, and subjected samples to storage at 4°C or 24°C and for different time periods^31^. They found consistent biological change with storage time and condition, but samples could still be assigned to the correct origin with supervised classification (random forest) among the studied sites. It is, however, still unclear how storage of soils impact basic biodiversity measures and compositional signatures of the original sampling site and habitat type in a broader ecological context.

Here, we quantify the changes in taxonomic composition for a soil sample, which was divided into multiple subsamples and stored under a range of conditions, with focus on storage time, temperature and exposure (in closed containers or in open containers allowing sample desiccation). We assessed biodiversity using eDNA metabarcoding targeting bacteria, fungi, protists (cercozoans), general eukaryotes and plants by use of taxon-specific primers (**Fig. 1**).

**Fig. 1.**
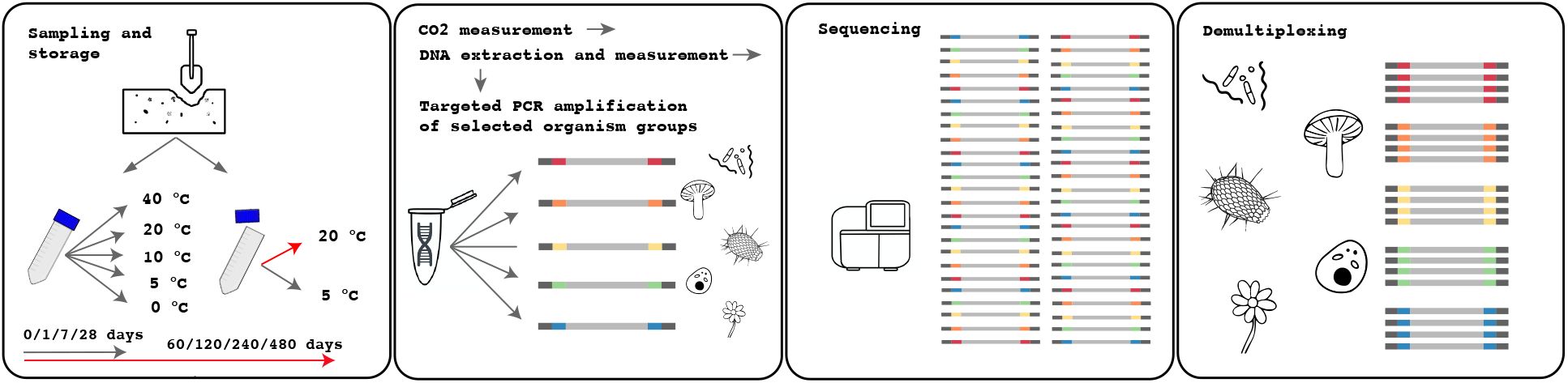
Experimental design. Soil samples were stored at different temperatures and exposures, and eDNA metabarcoding was performed targeting several organism groups. Soil was sampled, sieved and mixed/homogenized, and triplicates of tubes were subjected to storage at different combinations of temperatures, exposure, and storage time. Unexposed samples (tubes with lid on) were stored at 5°C, 10°C, 20°C and 40°C, and exposed soil (no lid) were stored at 5°C and 20°C. Tubes were harvested and analyzed after 0, 1, 7 and 28 days for all combinations of temperature and exposure. Tubes stored at 20°C (exposed and unexposed) were further harvested after 60,120, 240 and 480 days. 36 combinations of temperature, exposure and storage time were analyzed for a total of 108 samples. Analysis included respiration (CO2) measurement, DNA extraction and measurement and PCR amplification and sequencing (metabarcoding) of selected organism groups.

Our overall study aim was to assess the effect of sample storage on derived biodiversity metrics with a focus on biodiversity assessment and forensic applications. We approached this objective by investigating the following questions:

1) How does sample storage affect basic biotic patterns such as richness, evenness and taxonomic composition?

2) To what degree does sample storage change the signature of the sampling location and reduce the possibility of inferring the exact site of origin – i.e. sample matching?

3) To what degree does sample storage change the wider ecological signature and reduce the possibility of inferring the habitat type of the sampling site – i.e. sample provenance prediction?

Overall, we expected to see more change with longer storage time and higher storage temperature. Further, we expected to see most change for groups able to grow within our experimental systems, in particular bacteria and fungal molds, and least change for organisms directly relying on photosynthetic products, such as mycorrhizal fungi. We addressed point 1 by looking for significant changes in basic biodiversity measures, such as sample richness and evenness, community dissimilarity, as well as OTU change and change in taxonomic composition. To address points 2 and 3, we applied supervised learning. We employed a reference dataset, which covers all major terrestrial habitat types in the study area (Denmark). We chose to use k-nearest neighbors (KNN) as a simple supervised classification approach applied directly on overall community dissimilarity measures, as we did not aim for results directly dependent on presence/changes of particular taxa, and were interested in seeing the effect of storage on the full community.

## 3 Results

### 3.1 Absolute sample change from time zero with storage

Overall, measures of richness, evenness and community composition were relatively stable for all organism groups and systems ≤ 20°C for up to 28 days (**Fig. 2a-d, Supplementrary Table 2**), whereas measures diverged gradually for most systems stored at 40°C or stored at 20°C for 28 days or more. Generally, richness started to decrease after 1 to 28 days while evenness was more stable except for a few treatments. Community compositional dissimilarity to time zero started to increase at day 1 to 28. The concentration of DNA extracted was decreasing for 40°C samples from day 1, and for 20°C samples stored for more than 28 days (**Fig. 2e**). CO_2_ development per hour increased with storage temperature and decreased gradually with time for closed storage, whereas it was steady in open storage (**Fig. 2f**).

**Fig. 2.**
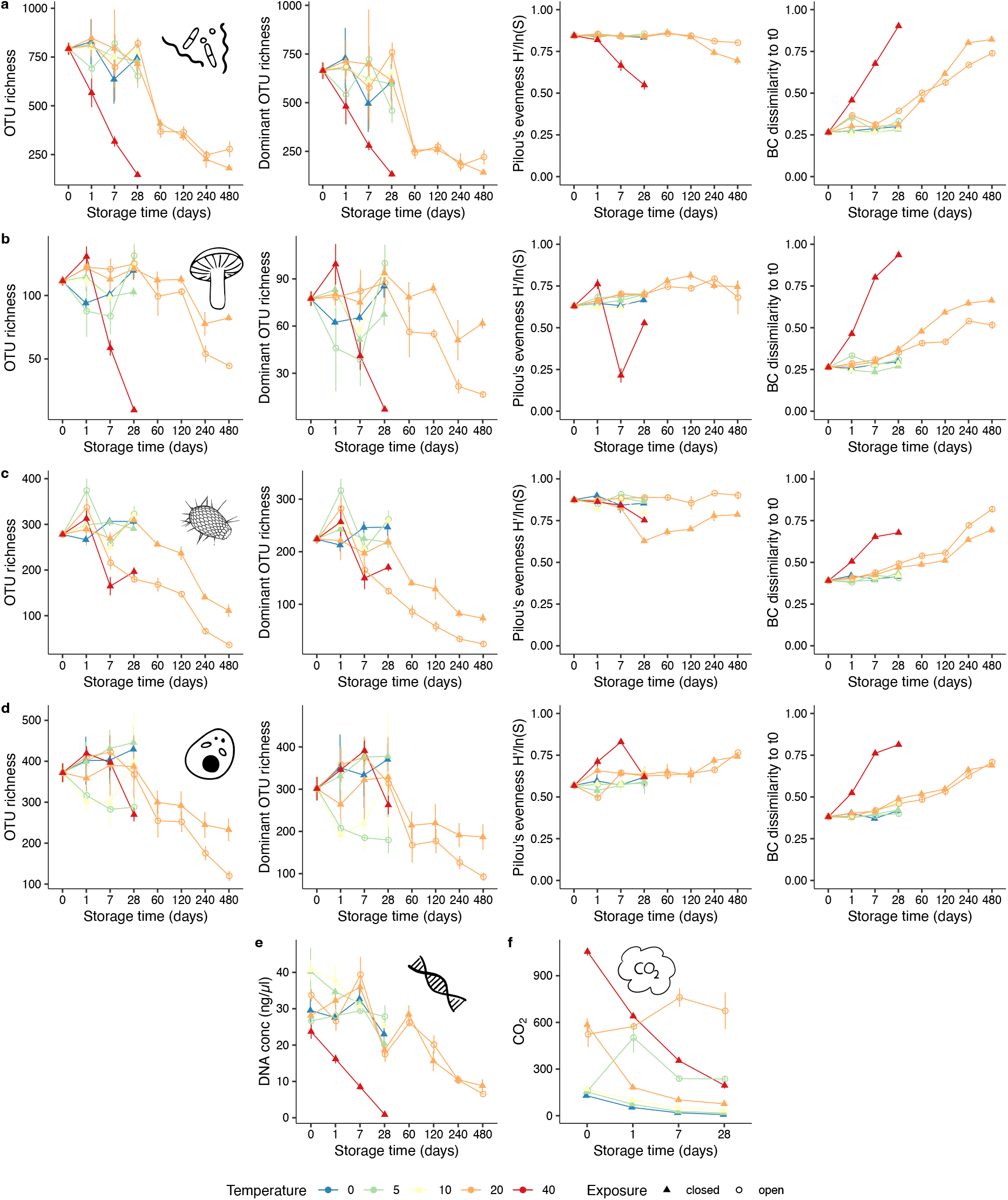
Absolute change with sample storage. Rows show (from top to bottom) a: bacteria, b: fungi, c: protists and d: eukaryotes, and bottom row shows, e: changes in measured DNA concentrations and f: measured CO2 development. Plots (a-d) show, column 1: change in OTU richness; column 2: richness of dominant OTUs (registered with ≥ 1% in each sample); column 3: community evenness as Pilou’s evenness index (H’/ln(S)); column 4: community change as Bray-Curtis dissimilarity from the centroid of the time zero communities. Plots show mean value +/- SEM for triplicates per treatment, with storage time on the x-axis, colors indicate storage temperature, and shape indicate exposure. Corresponding p-values for significant differences can be seen in Supplementary Table 2.

#### Richness (Fig. 2)

Generally, we found relatively large variation in richness estimates, and thus relatively few changes with time were significant though the trends were common for most taxa/treatment comparisons. The pattern for total richness and richness of dominant OTUs, were similar. Bacteria had the highest number of significant differences (20°C open & closed at 60 days or more, and 40°C at 7 days or more), whereas eukaryotes was the groups with fewest significant differences (40°C 28 days, and 20°C (open) at 240 days or more, and 5°C (open) at 7 days). Despite the lack of significance, the downward trend was evident for all 20°C samples stored for a long time, seemingly with a difference between open and closed tubes for fungi, protists and eukaryotes, where the open tubes showed a faster and more pronounced decrease in richness.

#### Evenness (Fig. 2)

The evenness of bacterial communities did not change significantly with time, although the figure shows a clear declining trend for 40°C (and partly 20°C closed) samples. For the other groups there were some significant differences, but generally, evenness was relatively stable with time. The protist data showed a marked difference for 20°C samples, where only the closed systems saw a drop in evenness from day 28.

#### Divergence from time 0 (Fig. 2)

All treatments gradually showed increased Bray-Curtis dissimilarity to time 0 community composition (the calculated centroid), with 40°C (and partly 20°C) samples increasing faster and more. All 40°C samples showed a clear and significant trend, being significantly more dissimilar from time zero already after 1 day. For most 20°C samples, divergence from time zero was apparent from day 28 day. For bacteria and fungi, the closed 20°C tubes changed faster and more than the corresponding open tubes, whereas protists showed the opposite pattern. The long term stored 20°C samples changed as much or more than the 40°C (28 day) samples for protists and bacteria. The results of the pairwise PERMANOVA (**Supplementary Table 3**) corresponded well with these finding, but only few adjusted p-values were << 0.05 due to the many comparisons.

#### Community change (Fig. 3)

In the NMDs ordinations of the communities, the samples stored at 0°C, 5°C and 10°C for up to 28 days, displayed no systematic change, reflecting the low level of change observed in the other metrics. However, for the samples stored at 20°C we observed a systematic change from day 28 and onwards, with open and closed tubes clearly showing different trajectories (least evident for fungi). The 40°C samples showed a clearly changed position already after 1 day of storage, and a different trajectory compared to the 20°C samples. For samples exhibiting evident change (20°C for 28 or more days and 40°C), the change was deterministic as triplicates generally remained close in the ordination (although the protists displayed some variation in the 20°C open samples at day 240 and 480).

**Figure 3.**
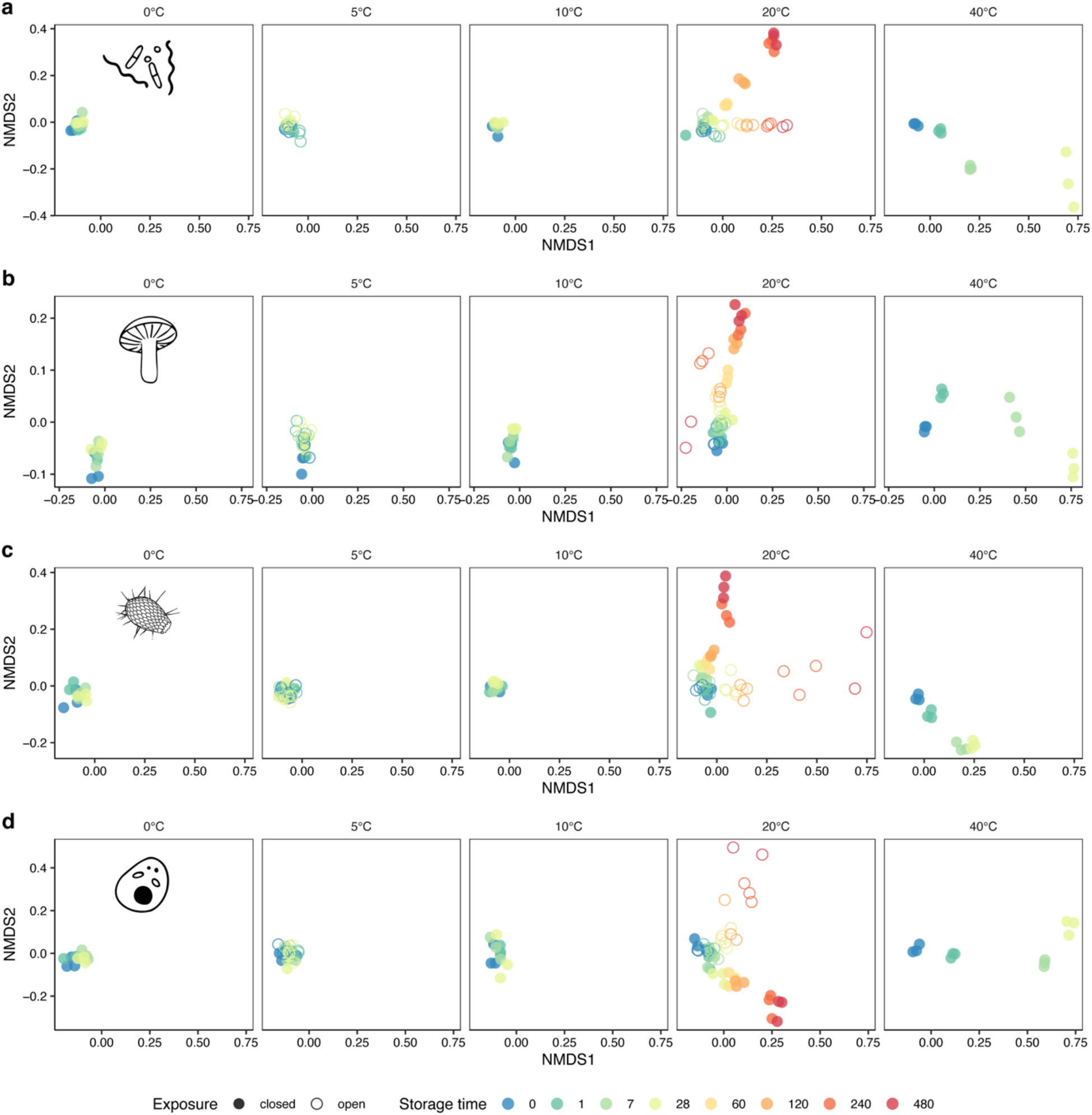
Community change with storage. NMDS ordinations of the stored samples showing community change with storage conditions and time for the four organism groups (a = bacteria, b = fungi, c = protists, d = eukaryotes). For each of the four organism groups, one NMDS ordination in 2 dimensions was performed on Hellinger transformed data. Axes show MNDS 1 and 2, colors indicate storage time, shape indicate exposure (open/closed tube), and facets reflect storage temperature.

**OTU change (Fig. 4)** was calculated by comparing the OTU composition of the combined triplicates of any treatment with the 21 combined time-zero samples. The expected OTU change due to stochasticity (without any storage effect) was 20 - 30 % corresponding to 2.6 - 5.9 % of the reads (**Supplementary Table 4**). **Fig. 4a** shows that the change did not exceed the expected change for most samples stored at 20°C or lower, whereas the 40°C samples showed a higher proportion of new OTUs per treatment – most so for the eukaryotes. The contribution of new OTUs to the total read composition (**Fig. 4b**) generally followed the same pattern for most treatments, except for long term storage and 40°C sample. For samples stored up to 28 days (and at 20°C or lower), the contribution of new OTUs resembled the level expected due to stochasticity. However, the bacterial data (20°C / closed) showed some late detected OTUs (day 28, 60 and 120) which later contributed a higher relative abundance than expected from stochasticity. This was also the case for fungi and eukaryotes, but less pronounced. For the protists, the relative contribution of new OTUs was generally low even for 40°C samples and was highest for 20°C open at 48 days, whereas the other three markers (bacteria, fungi and eukaryotes) by far showed the highest contribution of new OTUs in the 40°C samples already at day 7. The day 28 samples at 40°C for bacteria were mainly composed of reads of OTUs observed at day 7, whereas for fungi, OTUs observed already at day 1 dominated, and for eukaryotes they are dominated by equal amounts of OTUs observed at time zero and day 7.

**Fig. 4.**
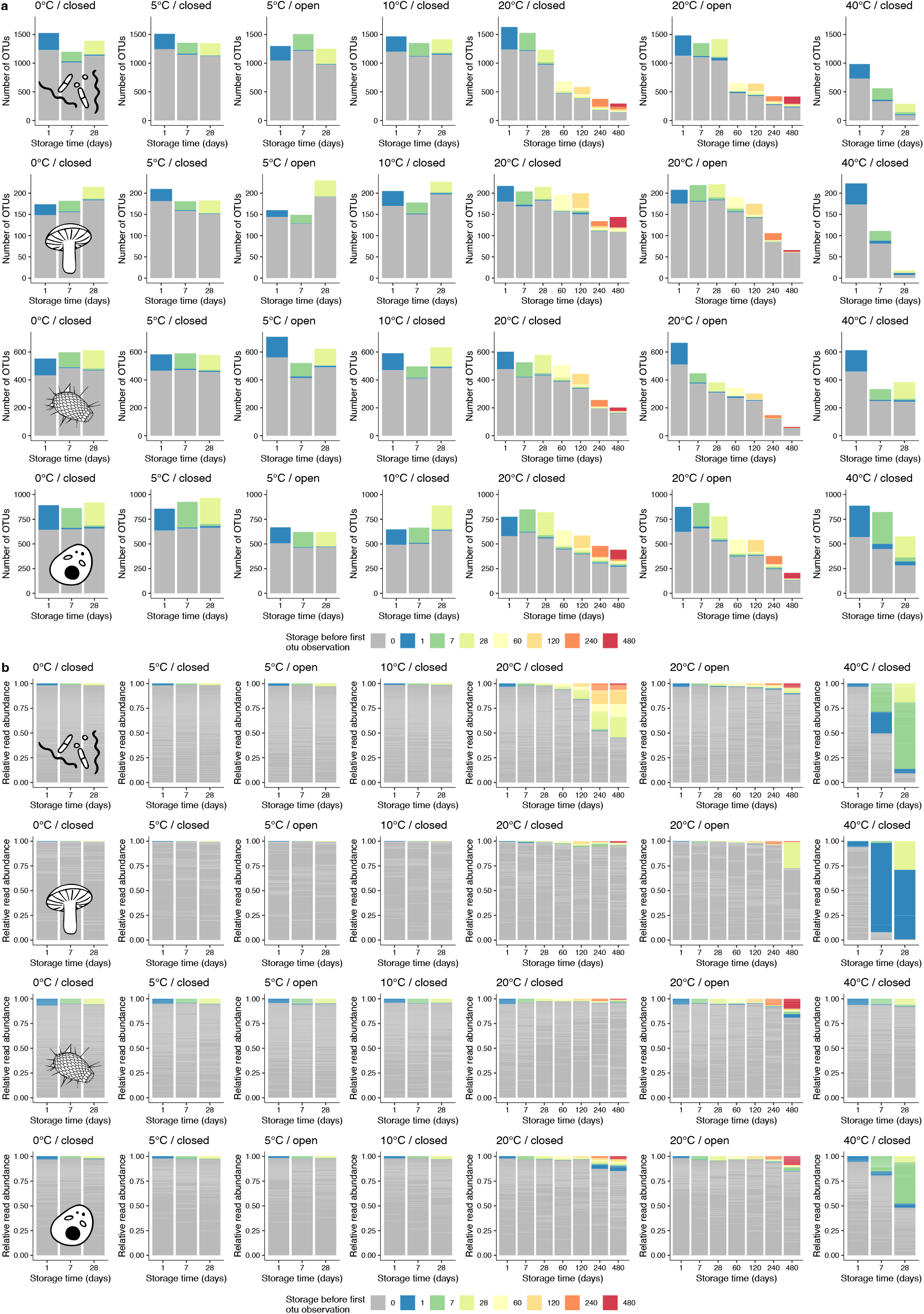
OTU change. Bar plots showing the OTU composition in terms of the first appearance for a given OTU. a) shows the composition of OTUs, b) relative read abundance of the OTU composition. Rows in each plot show (from top to bottom): bacteria, fungi, protists and eukaryotes. All three replicates of a given treatment (combination of storage time, temperature and exposure) are combined and compared to the combined composition of all 21 time-zero samples.

**Taxonomic changes (Fig. 5**, all taxonomic levels can be seen in **Supplementary Figs. 1-4**). ***Bacteria***: For most treatments of 20°C or lower, few major taxonomic changes occurred up to day 28. However, pronounced taxonomic changes took place in the 40°C samples where the Firmicute genus *Alicyclobacillus* increased to finally dominate the samples after 28 days. In the 20°C samples, gradual change in the proportions of several taxa was observed from day 60, and there was a clear difference between the open and closed tubes. The Firmicute genus *Bacillus* increased markedly after 120 days in the open tubes, whereas the closed tubes saw a corresponding increase of the Acidobacteria *Acidipila*. ***Fungi***: For all treatments of 20°C or lower, Mortierellomycetes (*Mortierella*) systematically increased, whereas Agaricomycetes (*Inocybe, Cortinarius*, etc.) concomitantly decreased already after 7 days. In the 40°C samples, *Aspergillus* dominated already after 7 days. ***Protists***: Reflecting the low OTU change, the taxonomic change of protists was less pronounced, but with a few systematic changes. The 20°C (open and closed tubes) displayed an increase of Allapsidae after 120 days, whereas, in the closed tubes, only Cryomonadida (Rhogostoma lineage) increased from day 7, and decreased again at day 240. Whereas the other organism groups displayed a drastically different taxonomic composition in 40°C samples, the taxonomic composition in the 40°C samples of the protists was comparable to that of samples at lower temperatures. ***Eukaryotes***: For most treatments of 20°C or lower at 28 days or less, few systematic taxonomic changes occured. For 20°C closed there was a decline of metazoan (mainly in the form of Enoplea nematodes) and an increase of fungi. This was also the case (but less linear) in the 20°C open tubes. This was also seen very clearly for the 40°C, where the Metazoa OTUs disappeared at day 7, and where an increase of Apicomplexa was also seen.

**Fig. 5.**
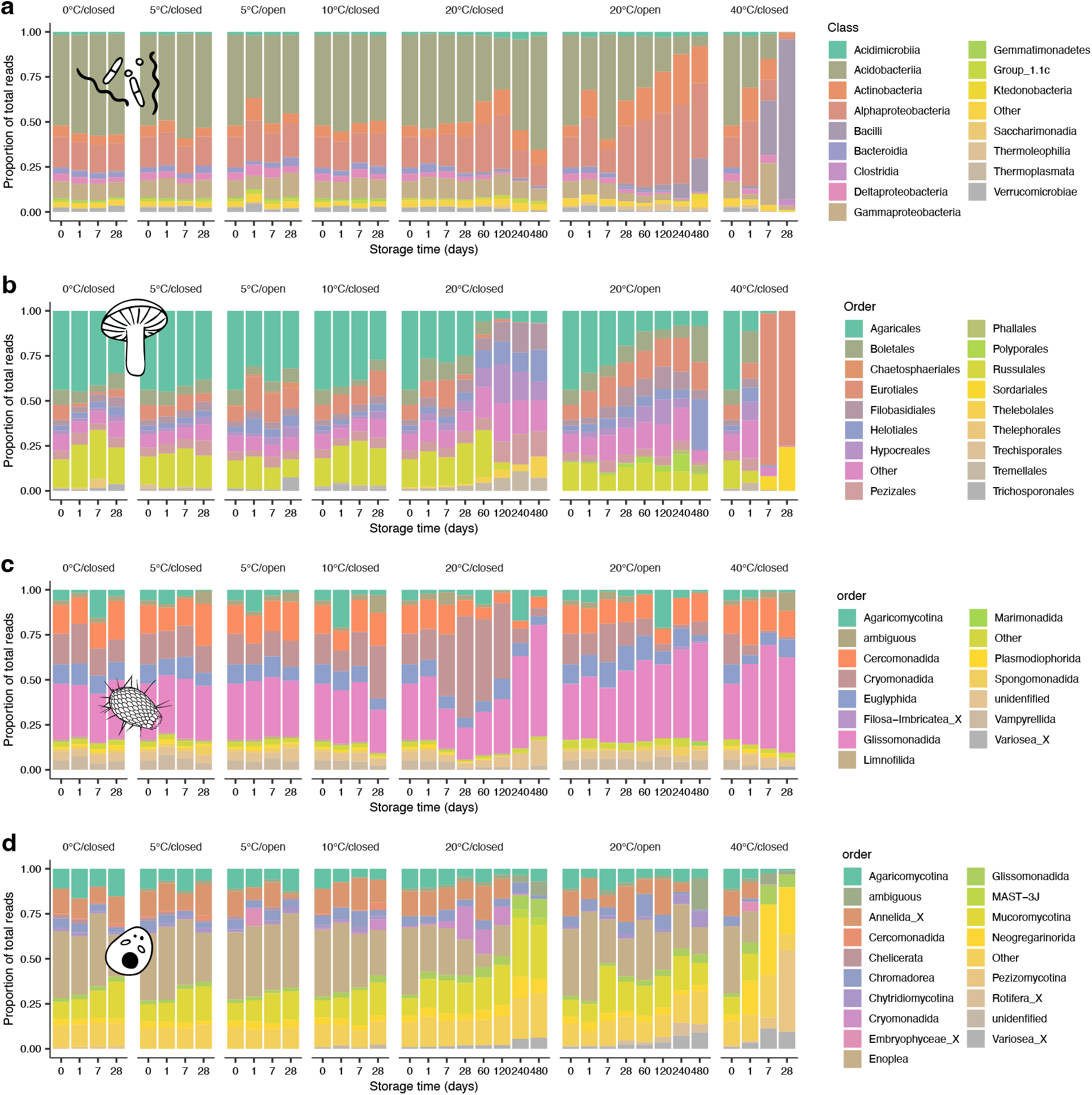
Taxonomic change with storage. Bar plot showing the relative composition of reads from the most abundant taxa at class level. Rows show bacteria (a), fungi (b), protists (c) and eukaryotes (d). All three replicates of a given treatment (combination of storage time, temperature and exposure) are combined. **Supplementary Figs. 1-4** show composition at all taxonomic levels.

### 3.2 Signature of exact location

We used supervised KNN classification to test if the stored samples could be reclassified to the correct location (sampling site) as represented by nine un-stored (time zero) samples using a 129 sample reference dataset as outgroup. Using a criterion of 0.5 mean probability, the approach classified all stored samples correctly (**Fig. 6 a-d**). The dissimilarity ratio – defined as the Bray Curtis dissimilarity of a stored sample to any of the 129 reference plots divided by the Bray Curtis dissimilarity to time zero centroid of the stored samples – became smaller with storage and temperature (**Fig. 7**), but the ratio never dropped below one. Thus, no sample changed to become more similar to other localities than to the origin. **Supplementary Fig. 5** shows how the absolute dissimilarity of stored samples to any of the 129 samples from the reference data show a steady increase for 40°C and long term (from day 28) storage 20°C samples.

**Fig. 6.**
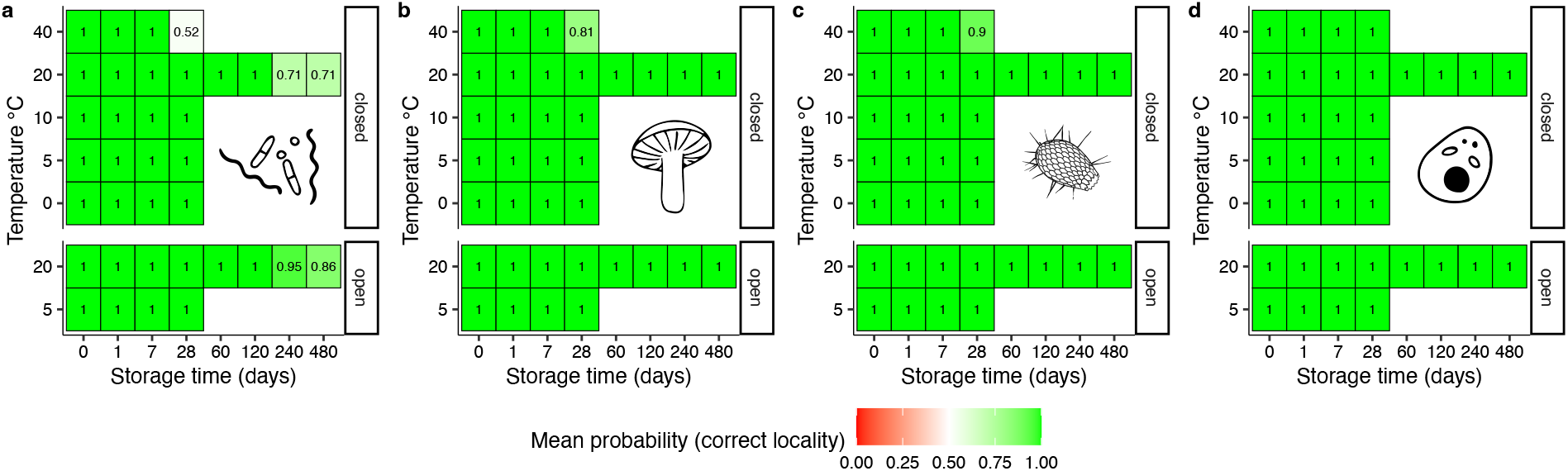
Supervised classification of exact location using k-nearest neighbor analysis (KNN). Probability of stored samples being classified as belonging to the exact sampling site, using nine time-zero stored samples as ingroup and 129 samples representing a wide selection of terrestrial habitats in Denmark as outgroup. Classification probability was calculated as the proportion of ingroup samples among the seven closest neighbors – defined as those samples with the smallest Bray-Curtis dissimilarity to the examined stored sample. Cells show the mean value of the triplicate per treatment. Panels show (from left to right): a: bacteria, b: fungi, c: protists and d: eukaryotes.

**Fig. 7.**
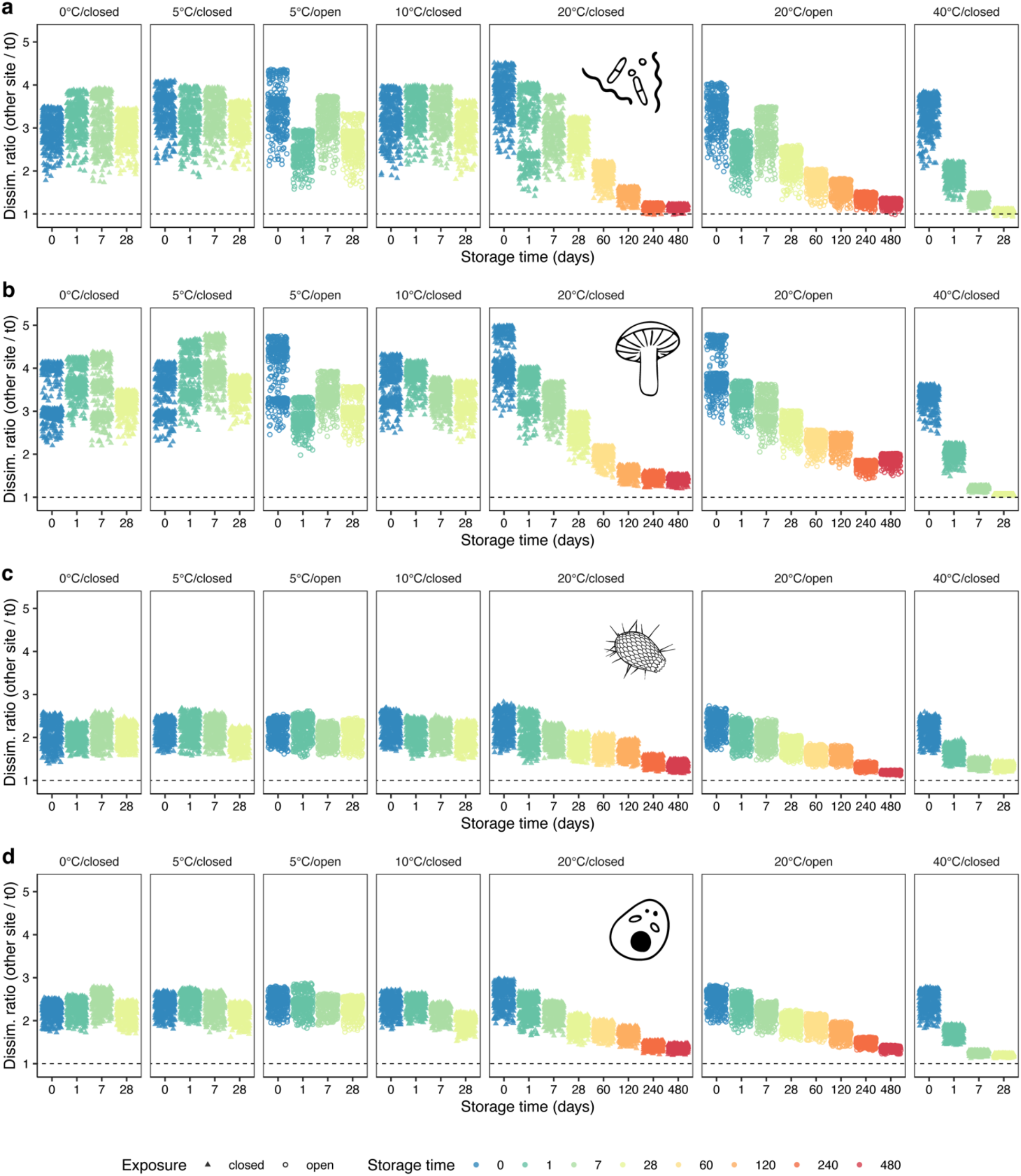
Dissimilarity ratio of stored samples to reference data compared to time zero. Each point shows the ratio between the Bray Curtis dissimilarity of a stored sample to one of the 129 reference plots divided by the Bray Curtis dissimilarity to time zero centroid of the stored samples. Thus, for each set of replicates for each treatment, the plot has 129 points. X-axis and color indicate storage time, symbol indicates exposure (open vs. closed tubes), and faceting corresponds to storage temperature and exposure. If a point is below the dotted line, it means that the stored sample is more similar to a reference plot than to the time zero centroid. This is not the case for any comparisons, meaning that the stored samples all retain highest similarity (lowest dissimilarity) to the time zero centroid compared to all reference plots.

### 3.3 Signature of habitat type

Using a supervised classification of the set of 130 reference sites into nine broadly circumscribed habitat types (see supplementary information), we used KNN classification to examine to which habitat type the stored samples were assigned. For all datasets, the dominant habitat type for un-stored samples was Mor forest, followed by Mull forest. This assignment fitted well with the ecological properties of the focal soil sampling site (SN081), which is mature beech forest on relatively poor, but not strongly leached, till with a top-soil pH of 3.9. This slightly ambiguous classification as acidic Mor forest borderline to alkaline Mull forest was seen even at time zero (**Fig. 8**), and also for the original reference sample from the focal site (SN081 indicated as “Ref” in **Fig. 8**). Using a criterion of 0.5 mean probability, **Fig. 8 a-d** shows that this approach classified all stored samples correctly, except for bacteria at 40°C after 7 and 28 days, and at 20 °C open after 480 days, and for fungi at 40 °C after 28 days. The probability of correct assignment was generally constant with storage time and with temperature. Apparent decline in assignment success was only seen for the for 40°C samples and for the 20 °C bacteria samples. Although the KNN approach was overall successful, **Fig. 8 e-h** shows that several stored samples exceeded the dissimilarity to the habitat type centroid by more than two standard deviations (for the reference data members of the habitat type). This was most evident for the protists although this group showed the highest and most stable mean classification success. **Supplementary Fig. 6** shows the classification probabilities towards the second most probable habitat type (Mull forest).

**Fig. 8.**
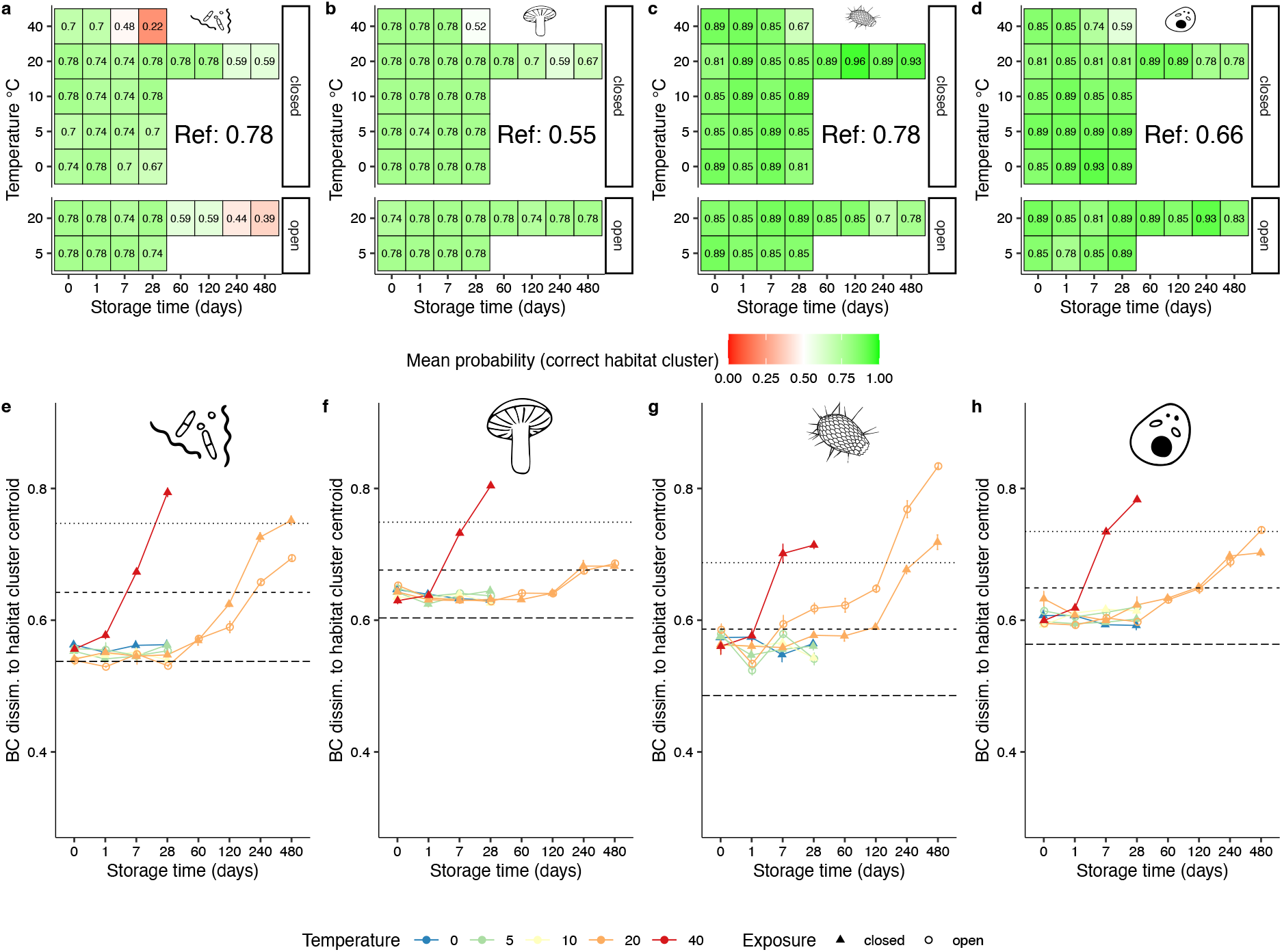
Supervised classification of habitat type and dissimilarity to habitat type centroid. *Upper panel* (a: bacteria, b: fungi, c: protists, d: eukaryotes) shows the probability of stored samples being classified as belonging to the correct habitat type among nine habitat types defined by supervised classification of observational data from the 129 reference sites, but without any samples representing the site of origin. Classification probability was calculated as the proportion of ingroup samples among the nine closest neighbors – defined as those samples (among the 129 reference samples only) with the smallest Bray-Curtis dissimilarity to the examined stored sample. Cells show the mean value of the triplicate per treatment. Note that only classification probability to the most probable habitat type (Mor forest) is shown. “Ref” indicates the classification success to the same habitat type of the origin site sample (SN081) from the reference dataset for comparison. *Lower panel* (e: bacteria, f: fungi, g: protists, h: eukaryotes) shows the Bray-Curtis dissimilarity of stored samples to the centroid of the most probable habitat type (Mor forest). Horizontal long-dashed line shows the mean dissimilarity to the centroid of the original habitat type members, the short-dashed line shows one standard deviation of the former, and the punctuated line shows two times standard deviation. Color indicates storage temperature and shape indicates exposure.

## Discussion

It is paramount in eDNA metabarcoding studies, that a sample adequately represents the community, from which it was drawn. Ideally, it should be comparable to an immediately processed sample, and only show deviations corresponding to what one would expect from the chosen analytical workflow. However, immediate processing is often not possible, and taxonomic compositional changes of the sample may occur. This is particularly the case in forensic sampling where storage conditions of soil traces are beyond the control of the analyst and for which storage under suboptimal conditions and for an unknown period of time is the norm. Depending on the objective of the study, some degree of community change during sample storage may be acceptable, but the uncertainties related to storage conditions are crucial to understand.

In this study, we addressed community change across a combination of storage conditions and periods. As expected, we found temperature-dependent community changes during storage time. However, we also observed that changes in measures of richness and diversity/evenness, and changes in community structure and taxonomic composition, were small for storage temperatures of 10°C (to 20°C) or lower and storage times of 28 days or less. However, biodiversity measures and community patterns diverged gradually for treatments at 40°C already after 1 day and for samples stored at 20°C for 28 days or more. Still, despite significant taxonomic compositional changes, we could still refer most samples to original habitat type and exact location with supervised classification models.

Our results show limited taxonomic compositional change during short-term storage (a few days) of samples. The variation in richness, evenness, community change and OTU change of samples stored at 20°C or lower and for 28 days or less did not exceed the level of variation for immediately analyzed (un-stored) samples. Thus, for studies of major biodiversity patterns, soil samples can be collected and stored for shorter time periods (days) without the need of immediate freezing/cooling, as long as it is possible to store samples at 20°C or lower. This will often be possible in temperate regions, and is practical if no lab facilities are nearby and/or if working with bulk samples, that need to be transported for further processing, etc. Still, targeting of certain fast-growing taxa, e.g. molds like *Mortierella* requires special consideration. On the other hand, our results show that higher temperatures (40°C) induce relatively early changes in taxonomic composition, as well as significant changes in other biodiversity measures already after one day. Hence, work in the tropics need special attention when there is not access to cooling. Desiccation is a good approach to conserve DNA^30,32^ and has also been used in practice for soil DNA studies with a global scope^6^. In this study, we only investigated passive desiccation in the form of open 20°C (and 5°C) tubes, which clearly differed from their closed 20°C counterparts, and the differentiation between closed and open treatments continued until the last sampling time. Whether this continuous change was due to differential growth of species present from the start, or partially from influx of new species to the open tubes is not clear. We expect that active desiccation with e.g. silica gel followed by storage in closed container may be the best approach, when cooling is not possible, as suggested by another study^30^. We saw a more or less identical pattern for total richness and richness of dominant OTUs, whereas other studies^30^ saw a marked difference with richness of dominant OTUs less sensitive to storage.

In this study, we combined the stored samples with a reference dataset representing most major terrestrial habitat types in Denmark including one sample from the same study site as the stored samples. Despite many differences in the sampling strategies of the two datasets, we could classify the stored sample to the correct habitat type using supervised classification in the form of simple KNN models.

The most important forensic lessons from this study, is that no stored sample gained higher similarity to any other sample after storage. Thus, all samples retained highest similarity to the original un-stored sample (time zero centroid) when compared to a reference dataset of 129 samples representing most terrestrial habitats in Denmark. Hence, the KNN models depending on compositional similarity could correctly match all stored samples to the correct exact location (as defined by the un-stored samples).

In forensic matching of samples – i.e. comparison of a trace sample to a crime scene – it is not permissible to get false matches, which may potentially lead to conviction of innocent persons. Thus, when employing community compositional approaches like this study, it is important to consider the strengths and weaknesses of the analytical approaches. The KNN approach uses dissimilarities to known observations, so if case evidence samples are merely investigated in the context of few other observations – or with observations from entirely different habitat types – false positives are likely. We suggest that real life forensic cases should not exclusively rely on approaches like KNN based on closest match, but also consider whether the observed dissimilarities lie within or close to the variation seen for replicated samples from the same locality, and ideally be combined with a score-based likelihood ratio-like measure. The matching approach applied here depends on a representative sampling with several replicates of the reference site. In the case of matching of two trace samples, or one trace sample with several compositionally diverse references, other approaches than KNN are needed.

Contrary to models for matching of forensic samples, models for provenance prediction (as defined and applied here) will most likely only be used as an investigative tool in forensic cases – e.g. to narrow down areas of interest – and thus some flexibility of models may be allowed as avoidance of false positive predictions is less critical. Here we tested whether the stored soil samples could correctly be classified to a wider habitat type of the location where they were collected. The KNN models for all organism groups were very successful and only failed for 40°C after 7 and 28 days for bacteria, and after 28 days for fungi.

For real-life forensic applications, we recommend prioritizing large representative ecological reference databases – i.e. sequence data from soils of a wide selection of habitat types – to reduce uncertainty in ecological inferences and site matching. Further studies are needed to test if such ecological reference database should be based on single bulk samples constructed for maximal representation of larger localities (habitat types) as used in this study, or one based on several replicates of smaller soil samples representing smaller or larger localities. Along with this, other sources of variation (like seasonality) also need to be addressed in future studies.

The changes we see in the stored samples are systematic – i.e. the replicates change in the same direction, as also detected in other in another study^33^. Furthermore, we see that the direction of community change depends on temperature and exposure (open/closed tube). It may thus be possible to predict storage condition and time for a questioned sample. This could be a valuable approach for forensic samples, where time since removal from the original site may be of interest, parallel to the estimation of post-mortem intervals. We also see clear differences in the taxonomic composition related to temperature and exposure, and it may be possible and interesting to identify indicator taxa for storage conditions. On the other side of this coin, it may also be possible to identify and extract those taxa that are least sensitive to storage and use these to build provenance and matching models that are robust irrespective of storage conditions. However, rigid examination of these topics would require soils from several different habitat types, as patterns likely depend on soil type.

In conclusion, this study shows that soil samples retains a large proportion of the original taxonomic compositional signature during relatively extended storage, and that the observed deviation – although deterministic – does not exceed the variance between replicated un-stored samples, if they are not stored warm or for a very long time. Still, this source of variation in biodiversity patterns from soil eDNA metabarcoding needs to be compared to other sources like seasonality, samples size, etc., to inform sampling strategies for biodiversity studies as well as making a solid foundation for interpretation of forensic analyses.

## 2. Materials and Methods

### 2.1 Experimental setup

We sampled soil in a mature beech (*Fagus sylvatica*) forest at the Strødam nature reserve in N Zealand, Denmark on August 31, 2017. The soil had a pH of 3.9 (H_2_O), a water content of 25% and organic matter content of 10%. Loose and coarse litter was removed from the soil surface before soil sampling, and the upper 10 cm was sampled, which then included a thin, ≈1 cm organic layer O and the top of the A horizon. The soil sample was taken from a single pit, about 5 liters in total in the middle of a permanently marked plot (SN081) established during the Biowide project ^34^.

Immediately after sampling, soil was carried to the laboratory (30 min drive) and sieved (5-mm mesh). 50-ml centrifuge (Falcon) tubes acted as experimental units and 3.2 g fresh weight of the sieved soil was added to each centrifuge tube. Tubes were then stored in combinations of temperature and exposure. The experimental setup was completed within 2-3 hours after field sampling (**Fig. 1**).

Five sets of tubes were closed with a lid to avoid desiccation and stored at 0°C, 5°C, 10°C, 20°C and 40°C, respectively. Two sets of tubes were left open to allow desiccation and stored at 5°C and 20°C, respectively. Tubes were harvested after 0 days (1 hour), 1 day, 1 week (7 days) and 4 weeks (28 days) and, further for tubes incubated at 20°C, after 2, 4, 8 and 16 months (60, 120, 240 and 480 days). All 36 treatments (experimental combinations of storage time, temperature and exposure) were in triplicate (i.e, n=108). Prior to storage, an 8-mm hole had been drilled into the lids and fitted with a rubber plug to allow for subsequent gas measurements. We used production of CO_2_ over time as a measure of total biological activity in our tubes. At each harvest event, CO_2_ production was measured for all tubes. The un-capped tubes were fitted with lids 30 mins before measuring CO_2_. After gas measurement, the harvested tubes were placed at -80°C for later DNA extraction and sequencing, while all remaining tubes were placed back at their respective incubation temperatures.

### 2.2 Measuring of CO_2_

We sampled gas from the headspace air from each sample tube with a gas-tight syringe inserted through the rubber plug. The 0.5 ml air sample was injected into a gas chromatograph equipped with a thermal conductivity detector (Mikrolaboratoriet, Århus) for the determination of CO_2_ concentration. Gases were separated before detection on a 1.8-m Haysep Q column operated at 45 °C. During each CO_2_ measuring event, we measured the CO_2_ concentration of atmospheric air and CO_2_ standards as appropriate.

### 2.3 Sequence data

#### 2.3.1 DNA extraction

DNA was extracted from 107 soil samples (originally 108 samples, but one failed) in two batches, one with the 84 tubes stored for 0, 1, 7 and 28 days, and the other with the 23 tubes stored for 60, 120, 240 and 480 days. From each sample, 0.25 g of soil was subjected to DNA extraction using DNeasy PowerSoil Kit (Qiagen), following the manufacturer’s protocol, except for the elution step, where 105 µl 1 x TET-buffer was used. For contamination control, five extraction blanks were included. Prior to extraction, the samples were homogenized using a TissueLyser II at 30 Hz for 10 min. DNA concentrations were measured with Qubit dsDNA HS (High Sensitivity) Assay Kit (Invitrogen) and samples were normalized to a concentration of 1 ng/μl prior to PCR amplification.

#### 2.3.2 DNA amplification and sequencing

DNA was amplified using five different markers targeting bacteria, fungi, protists (Cercozoa), general eukaryotes and plants, respectively (see **Supplementary Table 1** for primer and PCR information). The reason to include both a general eukaryote marker and specifically address fungi, plants and Cercozoa was to ensure appropriate amplification of some of our target groups, but still also to explore the usefulness of a more general primer but with less sequencing depth within specific clades. PCR reactions contained 0.04 U/μl AmpliTaq Gold (Life Technologies), 0.6 μM of each primer, 0.8 mg/ml bovine serum albumin (BSA), 1X Gold Buffer, 2.5 mM of MgCl, 0.2 mM of each dNTPs and 1 μl DNA extract in a 25 μl total reaction volume. Seven PCR blanks were included for every primer set. Fragment presence and sizes were verified on 2% agarose gel, stained with GelRedTM (Biotium).

Both forward and reverse primers were designed with 96 unique tags (MID/barcodes) of 6 bp at the 5′end using a restrictive dual indexing approach, where no primer tag (forward or reverse) was used more than once in any sequencing library, and no specific combinations of forward and reverse tag were reused in the study. PCR products were pooled for a total of 10 pools – two per primer set, with one pool containing the first 84 samples and another the remaining 23 samples. One or two extraction blanks, and two to three PCR negatives were included in each pool. PCR pools were purified with MinElute PCR purification kit (QIAGEN GmbH) and the length of PCR amplicons were verified on 2100 Bioanalyzer High Sensitivity Chip (Agilent Technologies). Each of the five pools containing 84 samples was built into four separate sequencing libraries, while pools containing 23 samples were built into one library per pool, four library negatives were also included (a total of 29 libraries). Libraries were built using the TruSeq DNA PCR Free Library Preparation Kit (Illumina), replacing all the manufacturer suggested clean-up steps (sample purification beads) with MinElute purification. A final library purification was carried out using Agencourt AMPure XP beads (Beckman Coulter) with 1.6 times beads to sample, and a final elution in 25 ul EB-buffer (Qiagen). Library concentration and presence of amplicons was verified with Qubit and BioAnalyzer (as above) and sequencing was done at the Danish National High Throughput DNA Sequencing Centre, on the Illumina Miseq v.3 platform (Illumina) with samples divided on four 300 bp paired end runs.

### 2.4 Post sequencing bioinformatic treatment

#### 2.4.1 Sequence processing

Bioinformatic steps followed the general procedures of earlier studies^35,36^ with minor modifications. Demultiplexing of samples was done with a custom script that keeps R1 and R2 separate for DADA2 processing, and is based on Cutadapt^37^ searching for a sequence pattern matching the full length combined tag and primer allowing for no errors, and removing possible remnants of the other primer at the 3’ end. We used DADA2 (v 1.8) ^38^ to identify OTUs as amplicon sequence variants (ASVs) and removal of chimeras (bismeras). For highly length variable markers (ITS2 for fungi), the script included a sliding window truncation of sequences from the 5’ end with Sickle^39^ (with options: pe -l 50 -q 28 -x -f -t sanger) to maximize output and quality of the ITS2 sequences that have length variation and therefore large differences in the onset of the quality drop towards the 3’ end. For the other markers where amplicon length is homogeneous, we applied a fixed length cutoff of the 5’ end, that allowed for ample overlap between R1 and R2 reads. Sequences were filtered and matched between R1 and R2 reads with DADA2 (using fastqPairedFilter with options maxN=0, maxEE=2, truncQ=2, matchIDs=TRUE). It has been advocated to use subsequent clustering and post-clustering curation to derive reliable biodiversity metrics^35^ (i.e. better species level OTU delimitation). However, this study was in part concerned with forensic application of environmental DNA, and we expected that intra-specific variation (artefacts or not) overall constitutes a reproducible signal, and therefore of potential value in forensic applications. Also, reproducibility and combinability of data is lowered by imposing arbitrary clustering levels and selection of representative sequences/centroids. Thus, we chose to apply our analyses to non-clustered OTUs (i.e. ASVs).

#### 2.4.2 Taxonomic assignment

For taxonomic assignment of OTUs, we used several different approaches. The bacterial data was assigned using the *assignTaxonomy* command in dada2 using the “silva_nr_v132_train_set.fa.gz” reference data. The fungal, protist and eukaryotic datasets were matched against reference databases using vsearch^40^ and a custom script that uses the top 10 matches to assign a majority rule taxonomy, and a similar approach was used for the plant data but using matches from BLASTn searches on GenBank. Assignment of the fungal data was done by matching the OTUs against the UNITE database for fungi^41^ and all eukaryotes^42^, and annotation of the Protozoa and eukaryote datasets was done by matching against the PR2 database^43^.

Forensic application would ideally utilize all data produced by a primer set to maximize reproducibility, whereas biodiversity studies generally work with focal taxonomic lineages. In this study, we only removed non-target sequences from the fungal dataset before downstream analyses, as these primers amplify a substantial amount of non-target (plant) sequences. For the plant sequence data, we could only identify six species of Viridiplantae – two vascular plants (*Fagus sylvatica* and *Hordeum vulgare*) and four green algae (*Bracteacoccus bullatus, Chlamydomonas hedleyi, Desmococcus olivaceus, Trebouxia decolorans*) – and this dataset was deemed too sparse to include in the remaining analyses.

### 2.5 Statistical analyses

For all analyses relying on OTU tables, the relevant table was resampled to the 25^th^ percentile to get even sequencing depth (but allowing a minor part of the samples to have lower read counts), as community dissimilarity measure, Bray-Curtis dissimilarity was used on Hellinger transformed OTU tables, and non-metric multidimensional scaling (NMDS) was done using the settings k=2, try=500, trymax= 4000 (using functions *rrarefy, decostand, vegdist* and *metaMDS* from vegan package^44^). All statistical analyses were run in R version 4.0.3 (2020-10-10) ^45^ on a x86_64-apple-darwin17.0 (64-bit) platform running under macOS Big Sur 10.16

#### 2.5.1 Absolute sample change from time zero with storage

Data from time zero samples (n=27, i.e. three from each of the nine combination of temperature and exposure) were used for the time zero population (reference) when analyzing effects of storage with time. To address changes in richness and diversity with storage, OTU richness was used as a proxy for total taxonomic/species richness. Following the findings of ^30^, we also measured the change in richness of dominant OTUs (OTUs registered with ≥ 1% in each sample). Pilou’s evenness index (H’/ln(S)) was used as a measure of evenness/diversity. To address change in community composition, we calculated Bray-Curtis dissimilarity between any stored sample and the centroid of all time zero samples. The centroid of time zero was calculated with the *dist_to_centroid* function (usedist package). For each particular treatment set (i.e. combination of temperature, exposure and storage time), we assessed significant changes in richness, evenness and community composition compared to the time zero communities, using t-tests with Bonferroni correction for multiple tests (i.e. 29 tests, excluding time zero combinations). Significant differences in community composition (compared to time 0) was also assessed with pairwise PERMANOVA as implemented in the function pairwise.adonis^46^ with the argument “reduce” to compare only against time 0). We considered p-values of <0.01 as significant. Community change was visualized with NMDS ordination.

Changes in OTUs over time was evaluated by identifying all OTUs observed for the first time at each storage time, using all OTUs from all 21 time zero replicates as a baseline of OTUs known to be present, and expected to potentially be detected after storage. Due to high microbial community complexity, sample heterogeneity and sampling stochasticity, we would expect OTU change between any sample comparisons. To calculate the expected number of new OTUs for any triplicate of (non-stored) samples, we randomly picked three of the 21 time zero samples and compared to the remaining 18 samples, 100 times. Furthermore, we evaluated the taxonomic composition for each treatment group, again combining triplicates per treatment.

#### 2.5.2 Relative change – habitat signature and forensic application

Despite of significant absolute changes in biodiversity metrics for stored samples, the change might still be insignificant for several applications, as the sample may have retained its signature – in terms of biological composition – of the exact sampling location or at least of the habitat type in a broader context. Therefore, to address the relative stability of the biotic signal of the stored samples, and thus the forensic utility and robustness of biodiversity measures, the stored samples were analyzed together with a reference dataset. The reference dataset stems from ^34^ and contains sequence data from 130 40m × 40m plots across Denmark. The 130 plots represent major gradients of moisture, fertility and succession, and thus include representatives of most natural to semi-natural habitats terrestrial habitat types in Denmark, as well as some agricultural and silvicultural land-use types ^34^. Soil samples from the reference dataset were collected and processed like the samples in this study, except that each of the 130 samples were constructed from a bulk sample of 81 smaller samples, that the soil was thoroughly mechanically homogenized (potentially releasing more intracellular DNA), that 4 grams of soil was used for the DNA extraction, and that the soil was sampled three years earlier in 2014 (November-December). The bulk sample used for the storage samples in this study was taken in the middle of one (SN081) of the plots used for the reference dataset, and this plot was excluded from those analyses where it could bias the interpretation.

Sequence data (OTU tables) from the present study and the reference dataset were combined for each of the four organism groups. Taxonomy was only assigned for the fungal data, to allow for exclusion of non-target sequences. For these combined analyses, we discarded OTUs with less than 10 reads in the reference dataset, and thereby excluded OTUs unique to the stored samples, that could otherwise make these samples more similar due to unique OTUs in that dataset. For supervised classification, we used k-nearest neighbor analysis (KNN) on community dissimilarity measures (Bray-Curtis dissimilarity of Hellinger transformed OTU tables).

##### 2.5.2.1 Signature of exact location

Using KNN, we investigated to which degree the stored samples retained characteristics of the exact location where they were collected, in the context of our reference dataset of terrestrial habitats in Denmark. Data from the reference dataset acted as outgroup. To avoid inflating classification success, we used only nine of the 27 time zero samples (triplicates of 0°C closed and 5°C closed and open) as ingroup. The soil used for the storage samples was sampled in the middle of one of the plots (SN081) from the reference dataset, so this sample could reasonably have been coded as ingroup. We chose, however, to exclude it from the models to not imposing any biases.

We calculated the proportion of ingroup and outgroup samples among the seven nearest neighbors (of the 129 reference plots and nine time zero samples) as the classification probability. As a direct visualization of the relative dissimilarities underlying this approach, we calculated and plotted a dissimilarity ratio for each stored sample in the form of the Bray-Curtis dissimilarities between the stored sample and the time zero centroid, compared to the dissimilarity between stored sample and each of the 129 reference plots.

##### 2.5.2.2 Signature of habitat type

Using a similar approach as above, we investigated to which degree the stored samples retained characteristics of the broader habitat type, to which the un-stored original sample was assigned. We used the survey dataset of 36 323 observations of 5 464 species (of vascular plants, bryophytes, macrofungi, lichens, and insects) recorded across the 130 reference sites (Brunbjerg et al 2019) to define nine strata (from hereon: habitat types), eight natural types and one agricultural. These habitat types were defined by supervised classification (see supplementary data) and encompassed the following: Mor forest, Mull forest, Bog forest, Swamp forest, Heathland, Grassland, Moor (acidic wetland), Fen (alkaline wetland), and Agriculture.

We then calculated the proportion of different natural strata among the nine nearest neighbors (of the 129 reference plots, excluding the reference sample from the sampling site of the stored samples, as well as all stored samples) as the classification probability. For comparison, we also calculated the classification probability for the original (SN081) reference sample from the sampling site. We established the variance of the ingroup stratum by calculating the Bray-Curtis dissimilarity of the cluster members to the habitat cluster centroid. Subsequently, the dissimilarity of stored samples to habitat centroids was related to the said variance in order to assess probability of correct habitat type assignment.

## Acknowledgments

This is a contribution to a national research project to develop soil forensic methods supported by Innovation Fund Denmark Grand Solutions (grant no. 6151-00002B, https://innovationsfonden.dk), and a contribution to a project on optimizing strategies for eDNA based biodiversity surveys, DNAmark, supported by Aage V. Jensen Foundation. The funders did not participate in study design, data collection and analysis, decision to publish, or preparation of the manuscript. Jens H. Petersen is thanked for designing four of the organism group icons.

## Author contributions

RK, TGF, FE, MV, AJH, RE conceived the overall study. RK carried out the sampling and designed the storage setup. IBN did the lab work. TGF, RK and RE decided on the analytical approach. TGF did the bioinformatics, statistical analyses, and plots. TGF and RK wrote the first draft. All authors contributed to the finalizing of the manuscript.

## Additional information

Supplementary Information accompanies this paper at DOIxxx

### Competing interests

The authors declare no competing financial interests.

The following Additional Information is available for this article:

## Supplementary Information 1. Construction of habitat types (clusters) for supervised classification

We investigated to what degree the stored samples retained characteristics of the same broader habitat type to which the un-stored original samples belonged. We aimed for a simple habitat classification that would be ecologically meaningful. A parallel aim was a relatively easy visual recognition of the resulting types, in order to ensure forensic applicability, i.e. that provenancing analyses would point to habitat types that can be identified by the police without compulsory assistance from ecological expertise. Thus, we chose to define these habitat types from major abiotic gradients (hydrology, soil pH/fertility, successional stage/vegetation structure) and re-classify it using species composition of the above-ground biota (not soil eDNA). These ecological complex gradients have proven by far the most important governing species composition of terrestrial communities of plants, animals and macrofungi^1^. Using standardized methods, site species data on vascular plants, bryophytes, macrofungi, lichens, gastropods and arthropods were collected from the same 130 study sites (40 × 40 m) as those we used in the eDNA analyses^2^. The survey data set contained 36 323 observations of 5 464 species recorded across the 130 reference sites. The original inventory was based on 25 design strata, representing the mentioned three complex gradients and a more detailed array of agricultural and silvicultural types. For the present study, we simplified these strata to eight natural types (combinations of canopy-covered vs. open, poor/acidic vs. rich/alkaline, and dry vs. wet) and one agricultural (rotational fields and lays): viz. Mor forest, Mull forest, Bog forest, Swamp forest, Heathland, Grassland, Moor (acidic wetland), Fen (alkaline wetland), and Agriculture. The discrimination in the original stratification between plantation forest and natural forest was abandoned in the simplified classification.

Using these nine strata, we applied supervised learning to adjust the classification in order to best reflect the actual above-ground species composition. We did an NMDS ordination in six dimensions (metaMDS, try = 100, trymax = 200), and used the first four dimensions for quadratic discriminant analysis. This resulting model was used to reclassify the 130 sites. Only three sites changed class assignment as a consequence of the re-classification, i.e. one Agricultural to Grassland, one Mor Forest to Mull forest and one Mull Forest to Mor forest. The focal site of soil sampling for the present study was borderline between Mull Forest to Mor forest.

**Supplementary Table 1.**
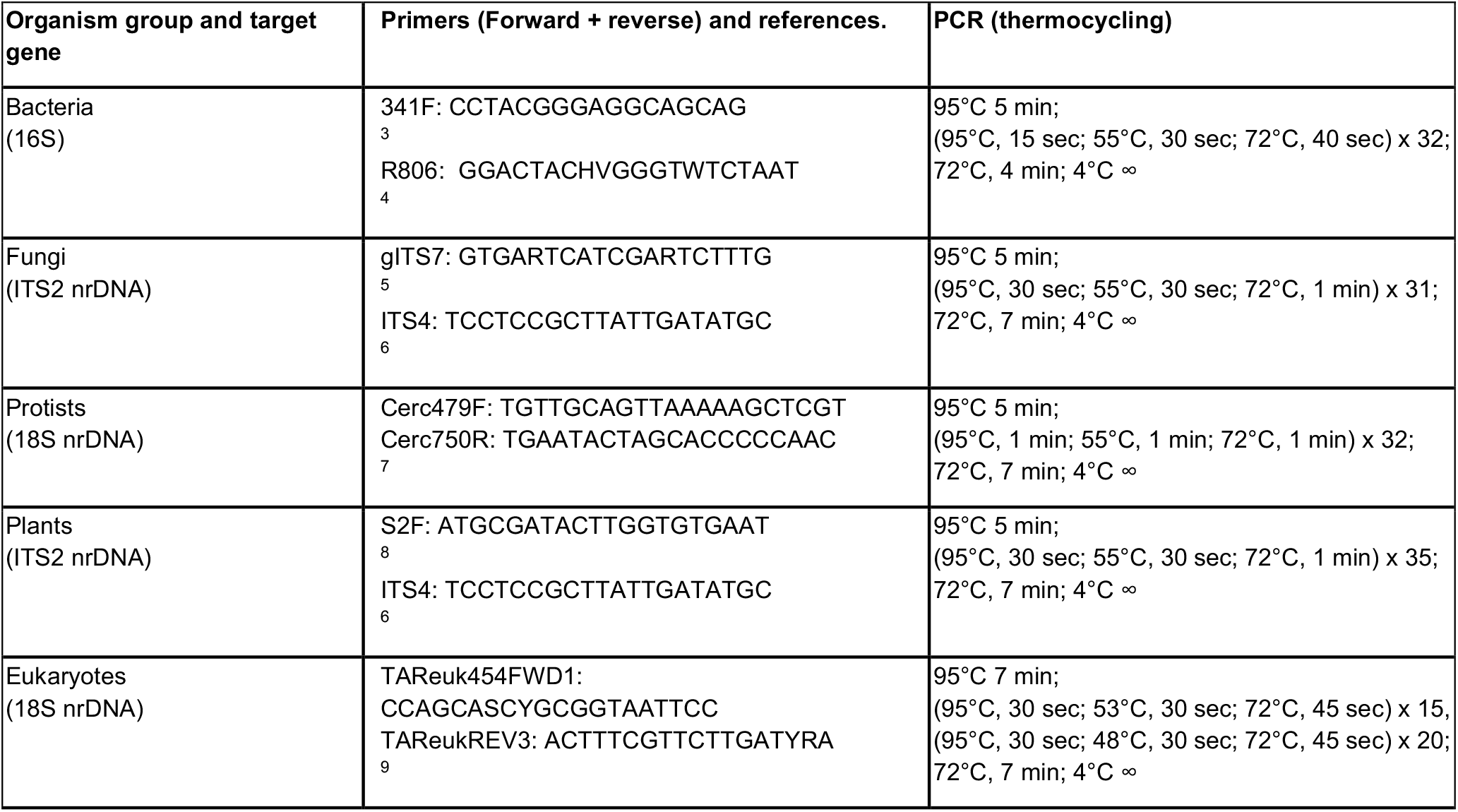
Information of primers and amplification.

**Supplementary Table 2.**
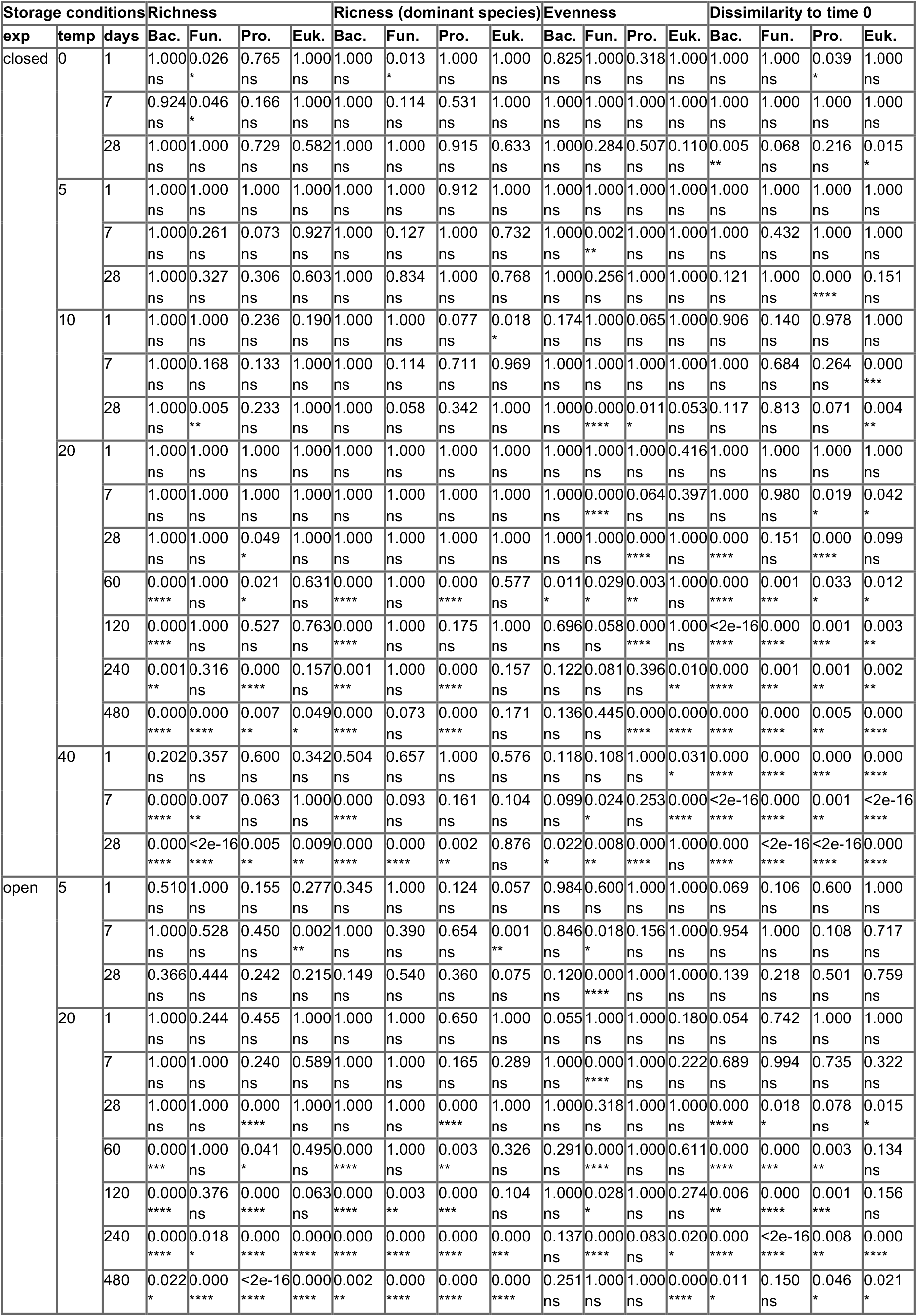
Changes in biodiversity measures with storage. Pairwise t-tests for significant difference in measures compared to time 0 (not stored) samples. All 29 time-0 samples were used as part of the time-0 population. P-values were corrected for multiple comparisons.

**Supplementary Table 3.** Community change with time. The table show adjusted p-values (Bonferroni) for the PERMANOVA tests of community change of stored samples against un-stored samples using the function pairwise.adonis with “reduce” argument to restrict comparisons with time zero (all 21 time zero samples acted as time zero reference).

| Exposure | Temperature | Storage time | Bacteria | Fungi | Protists | Eukaryotes |
| --- | --- | --- | --- | --- | --- | --- |
| closed | 0 | 1 | 1 | 1 | 1 | 1 |
| closed | 0 | 7 | 1 | 1 | 1 | 0.551 |
| closed | 0 | 28 | 1 | 1 | 1 | 1 |
| closed | 5 | 1 | 1 | 1 | 1 | 1 |
| closed | 5 | 7 | 1 | 1 | 1 | 1 |
| closed | 5 | 28 | 1 | 1 | 1 | 1 |
| closed | 10 | 1 | 1 | 1 | 1 | 1 |
| closed | 10 | 7 | 1 | 1 | 1 | 1 |
| closed | 10 | 28 | 1 | 0.464 | 1 | 0.029 * |
| closed | 20 | 1 | 1 | 1 | 1 | 1 |
| closed | 20 | 7 | 1 | 1 | 1 | 1 |
| closed | 20 | 28 | 0.116 | 0.029 * | 0.029 * | 0.087 . |
| closed | 20 | 60 | 0.029 * | 0.058 . | 0.029 * | 0.029 * |
| closed | 20 | 120 | 0.029 * | 0.029 * | 0.029 * | 0.029 * |
| closed | 20 | 240 | 0.058 . | 0.116 | 0.058 . | 0.058 . |
| closed | 20 | 480 | 0.087 . | 0.029 * | 0.029 * | 0.058 . |
| closed | 40 | 1 | 0.058 . | 0.058 . | 0.029 * | 0.029 * |
| closed | 40 | 7 | 0.029 * | 0.029 * | 0.029 * | 0.087 . |
| closed | 40 | 28 | 0.029 * | 0.029 * | 0.058 . | 0.058 . |
| open | 5 | 1 | 0.029 * | 1 | 1 | 1 |
| open | 5 | 7 | 1 | 1 | 1 | 1 |
| open | 5 | 28 | 0.435 | 1 | 1 | 1 |
| open | 20 | 1 | 0.029 * | 0.754 | 1 | 1 |
| open | 20 | 7 | 1 | 0.116 | 1 | 0.551 |
| open | 20 | 28 | 0.029 * | 0.029 * | 0.029 * | 0.058 . |
| open | 20 | 60 | 0.029 * | 0.058 . | 0.029 * | 0.058 . |
| open | 20 | 120 | 0.058 . | 0.029 * | 0.029 * | 0.029 * |
| open | 20 | 240 | 0.029 * | 0.058 . | 0.087 . | 0.029 * |
| open | 20 | 480 | 0.174 | 0.203 | 0.29 | 0.087 . |

**Supplementary Table 4.** Expected new OTUs in sample. Expected number of new OTUs when sampling 3 new replicates compared to the other 18 time zero replicates. The second column shows the expected number of new OTUs (+/- 1 sd) and the mean percentage of total OTUs this constitutes. The last column shows how large a proportion of the total reads these expected new OTUs represent. Values are estimated by randomly selecting 3 of the 21 time zero replicates and comparing the AOTU composition with the remaining 18 time zero replicates. The values are used to compare with the observed contribution of new OTUs of the stored samples compared to all 21 time zero samples.

| Organism group | # new otus | Relative read abundance of new OTUs (%) |
| --- | --- | --- |
| Bacteria | 334 ± 76 (22.1 %) | 2.6 ± 0.7 |
| Fungi | 42 ± 7 (19.8 %) | 3.2 ± 4.9 |
| Protists | 120 ± 13 (21.5 %) | 5.9 ± 0.8 |
| Eukaryotes | 258 ± 52 (30.2 %) | 3.7 ± 1.3 |

**Supplementary Figure 1.**
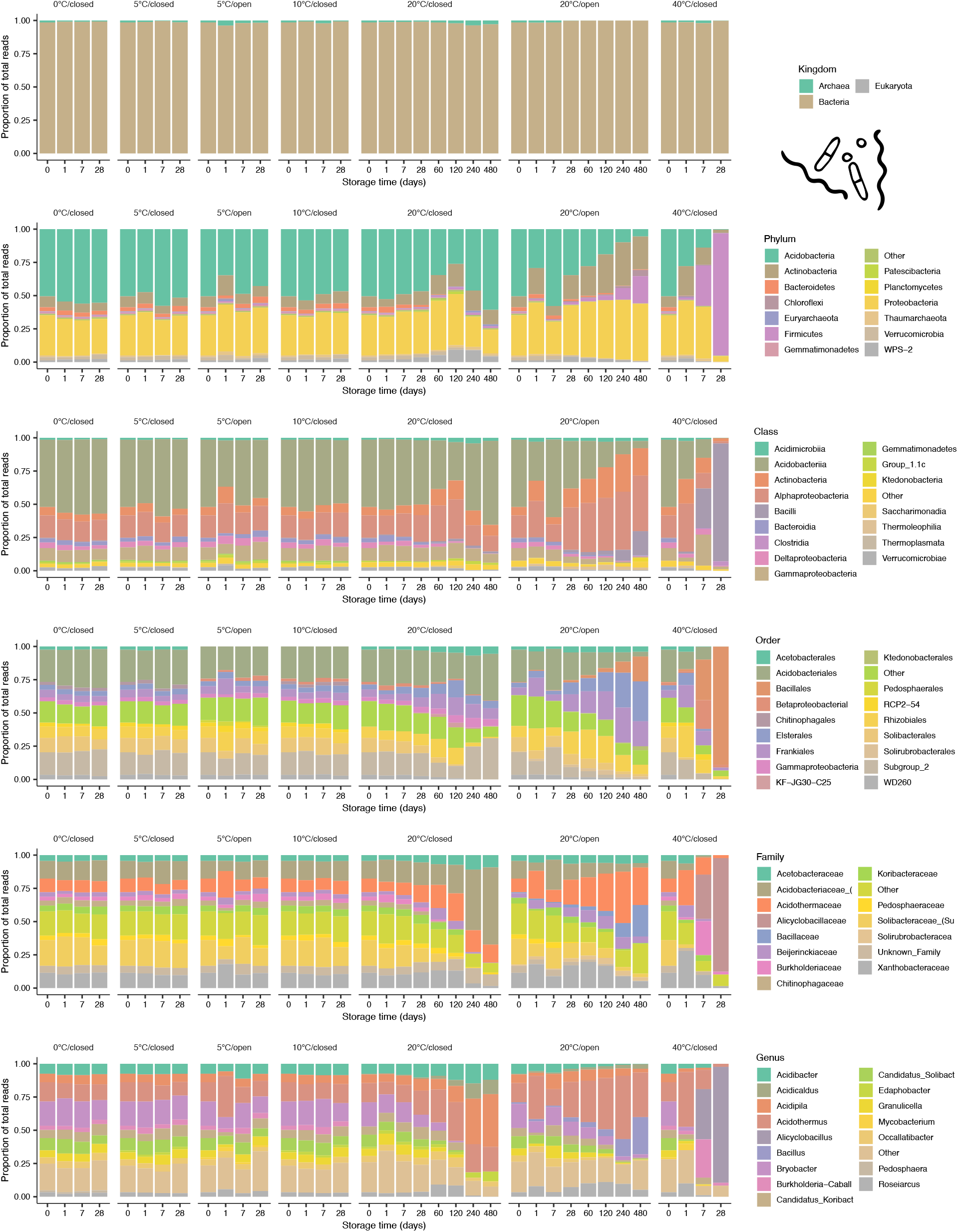
Taxonomic change with storage (bacteria). Bar plot showing the relative composition of reads from the most abundant taxa at major taxonomic levels for different combinations of storage conditions (temperature, exposure) and time. All three replicates of a given treatment (combination of storage time, temperature and exposure) are combined.

**Supplementary Figure 2.**
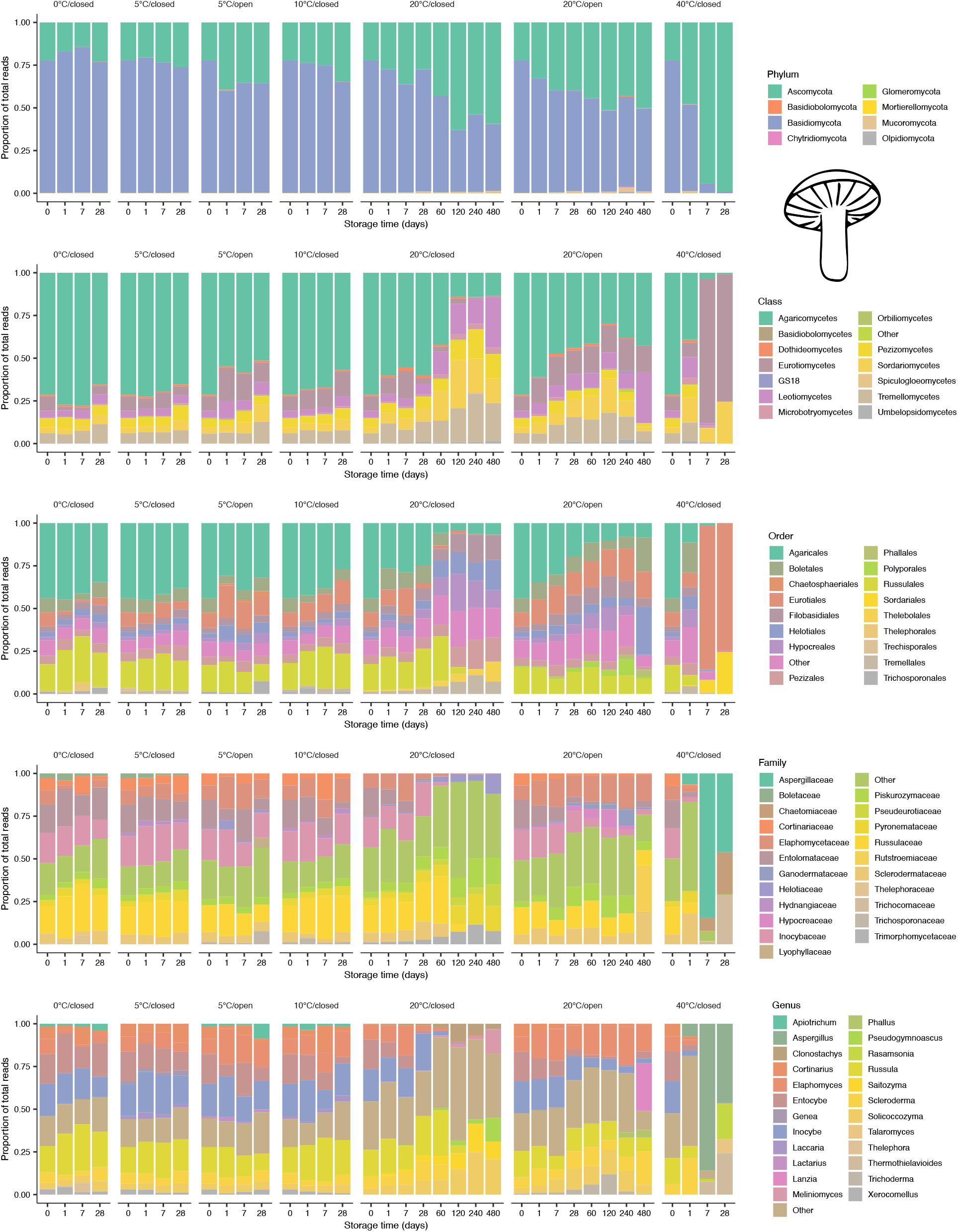
Taxonomic change with storage (fungi). Bar plot showing the relative composition of reads from the most abundant taxa at major taxonomic levels for different combinations of storage conditions (temperature, exposure) and time. All three replicates of a given treatment (combination of storage time, temperature and exposure) are combined. Kingdom level not shown (all reads not assigned to Fungi were removed).

**Supplementary Figure 3.**
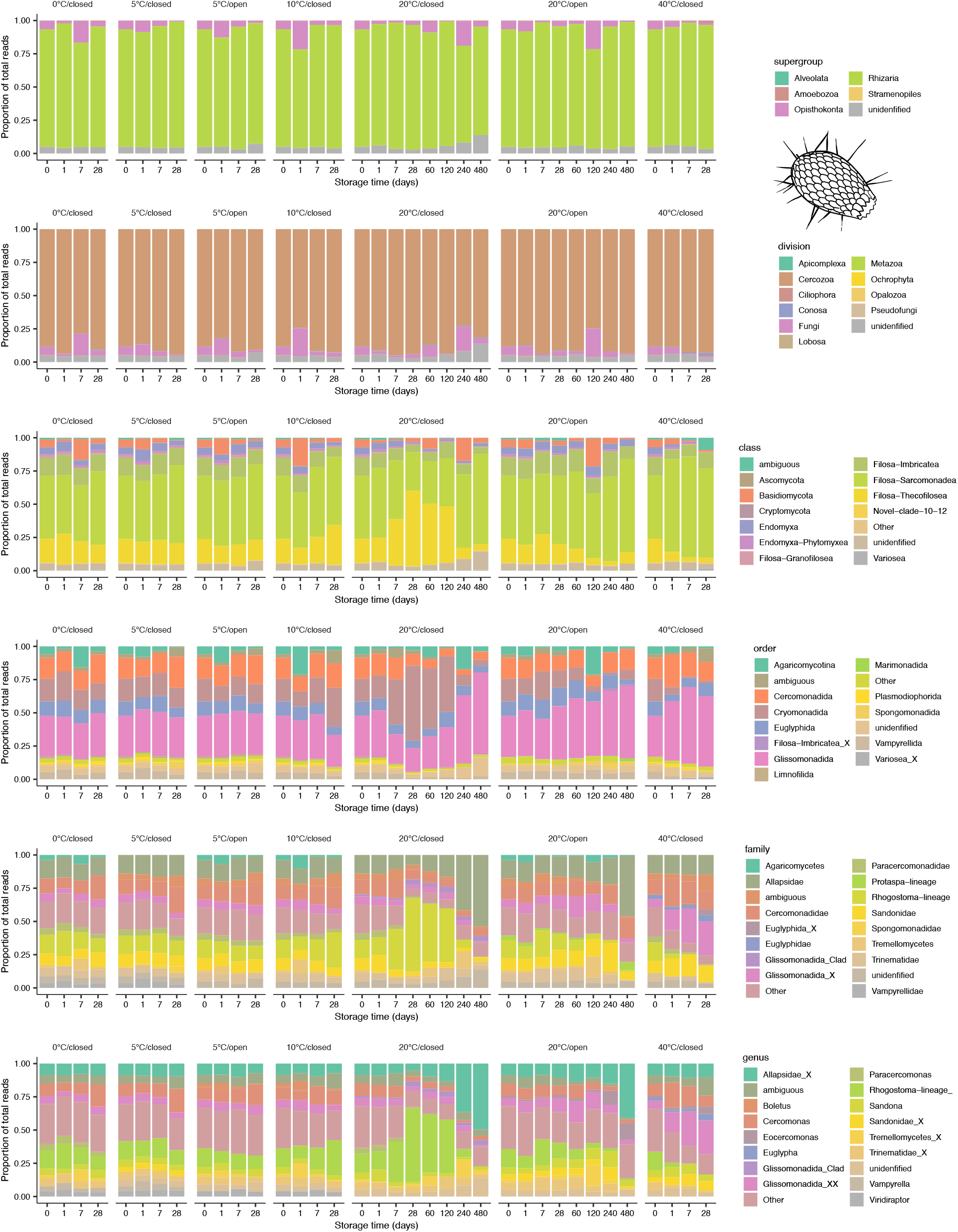
Taxonomic change with storage (protists). Bar plot showing the relative composition of reads from the most abundant taxa at major taxonomic levels for different combinations of storage conditions (temperature, exposure) and time. All three replicates of a given treatment (combination of storage time, temperature and exposure) are combined. Kingdom level not shown (all reads were assigned to Eukaryota except a few unassigned).

**Supplementary Figure 4.**
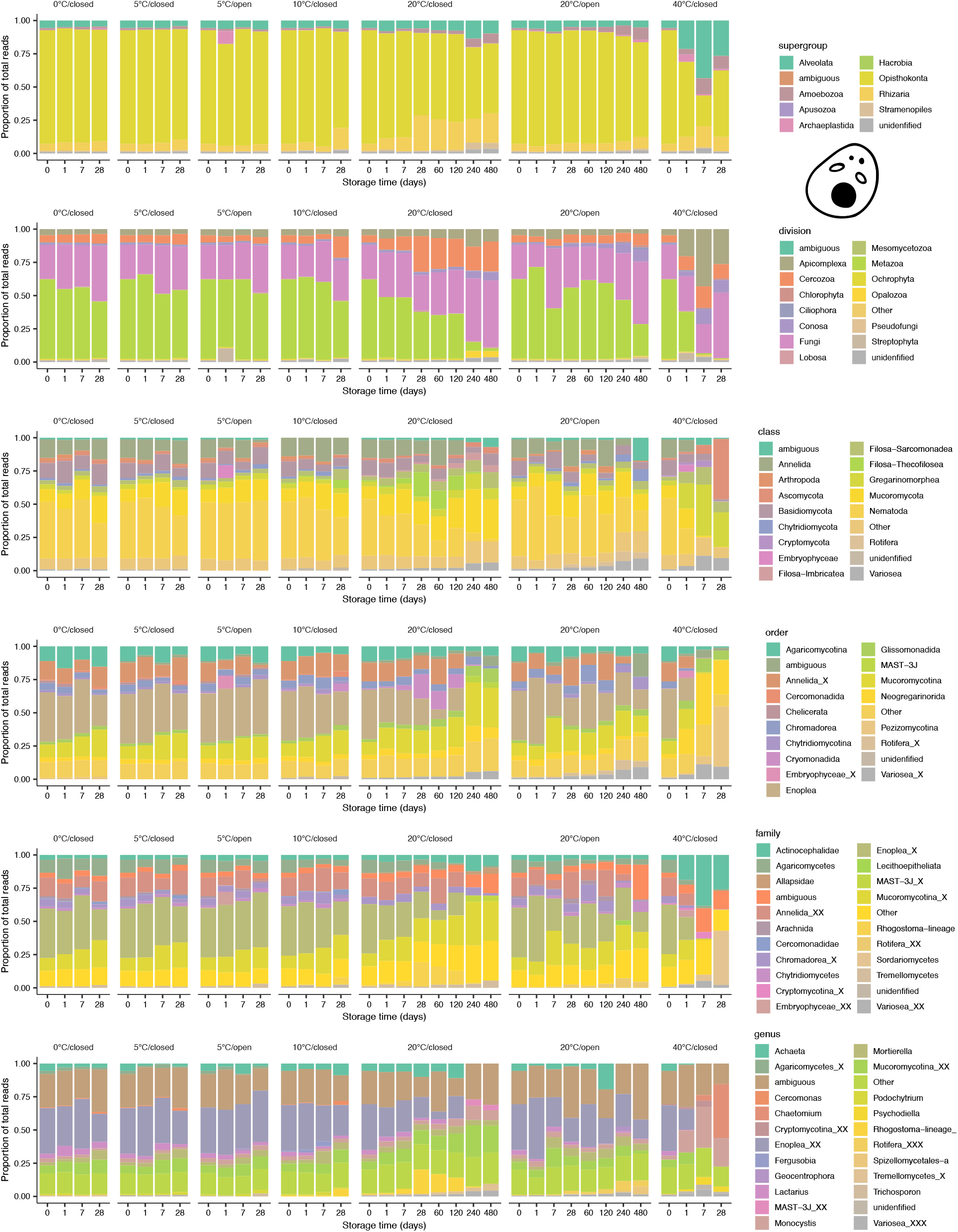
Taxonomic change with storage (eukaryotes). Bar plot showing the relative composition of reads from the most abundant taxa at major taxonomic levels for different combinations of storage conditions (temperature, exposure) and time. All three replicates of a given treatment (combination of storage time, temperature and exposure) are combined. Kingdom level not shown (all reads were assigned to Eukaryota except a few unassigned).

**Supplementary Figure 5.**
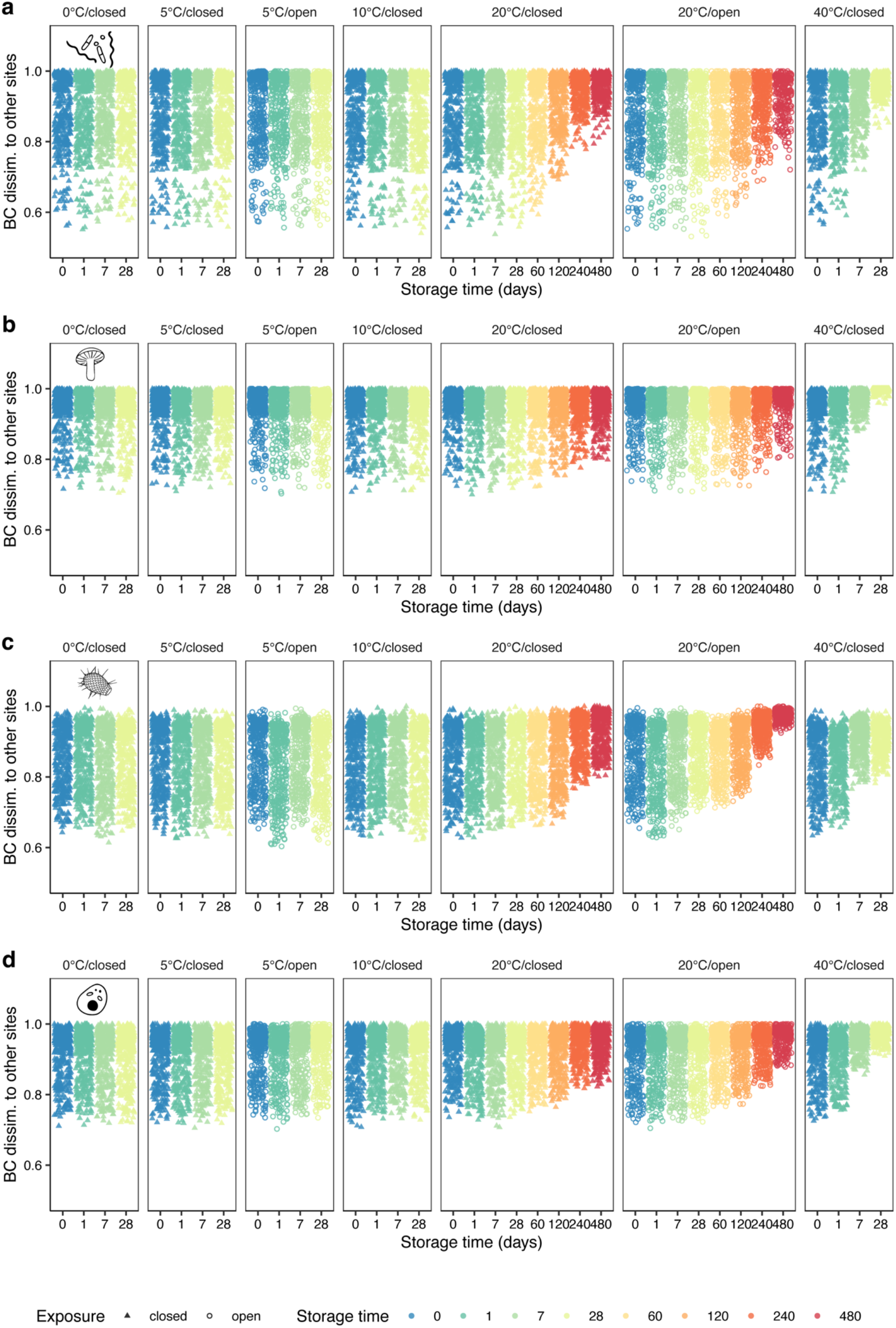
Community dissimilarity of stored samples to reference data. Each point shows the Bray-Curtis dissimilarity of a stored sample to one of the 129 reference plots (excluding SN081, which was the collected at the same locality as the stored samples). X-axis and color indicate storage time, shape exposure (open closed tubes), and faceting corresponds to storage temperature and exposure. The plots shows that storage does not result in increased similarity (decreased dissimilarity) to other sites. NB: Y-axis truncated at 0.5

**Supplementary Figure 6.**
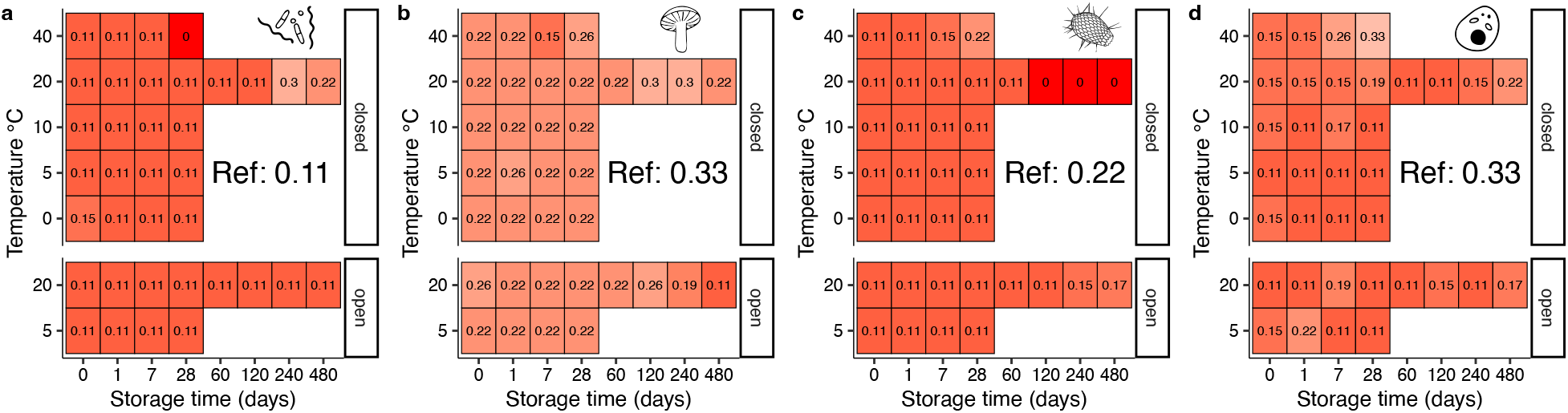
Supervised classification of habitat type and dissimilarity to habitat type centroid. *Upper panel* (a: bacteria, b: fungi, c: protists, d: eukaryotes) shows the probability of stored samples being classified as belonging to the second most probable habitat type among nine habitat types defined by supervised classification of observational data from the 129 reference sites, but without any samples representing the site of origin. Classification probability was calculated as the proportion of ingroup samples among the closest neighbors – defined as those samples (among the 129 reference samples only) with the smallest Bray-Curtis dissimilarity to the examined stored sample. Cells show the mean value of the triplicate per treatment. Note that only dissimilarity to the *second most probable* habitat type (Mull forest) is shown. “Ref” indicates the classification success to the same habitat type of the origin site sample (SN081) from the reference dataset for comparison.

